# The ER stress transcription factor Luman/CREB3 is a novel regulator of Schwann cell survival and myelinating capacity through the activation of the unfolded protein response and cholesterol biosynthesis pathways

**DOI:** 10.1101/2024.09.22.614000

**Authors:** Justin Michael A. Naniong, Vikram Misra, Valerie M. K. Verge

**Author notes:** Corresponding author: Valerie M. K. Verge, Cameco MS Neuroscience Research Centre, Saskatoon City Hospital, Saskatoon, SK S7K 0M7, Canada. Montreal Clinical Research Institute (IRCM), 110 Pine Avenue West, Montreal, Quebec H2W 1R7, Canada.

## Abstract

Misfolded protein accumulation in demyelinating disorders or injury can trigger internal endoplasmic reticulum (ER) stress alleviation mechanisms, namely the unfolded protein response (UPR) and cholesterol biosynthesis pathways. Here we demonstrate that the ER stress-associated transcription factor Luman/CREB3, herein called Luman, shown to drive axon regeneration by UPR-dependent regulation of adaptive low-level stress, also positively regulates rat Schwann cell (SC) survival, myelination, the UPR, and cholesterol production *in vitro*. siRNA knockdown of SC Luman expression decreased SC viability and increased apoptosis 48 hours post-transfection. Dorsal root ganglion (DRG) neuron/SC co-cultures where SCs overexpressed Luman exhibited increased myelination. Simulation of unstressed, mild and moderate ER stress by tunicamycin-mediated UPR induction in SCs, allowed examination of Luman on expression of known UPR regulators, including *Xbp1, Xbp1s*, CHOP, and pIRE1. They collectively demonstrate a cytoprotective role for Luman under manageable ER stress. Total cholesterol levels and sterol precursor *Srebf1* expression, key to myelination, also decreased following Luman knockdown. Finally, levels of mature brain-derived neurotrophic factor (mBDNF), a positive regulator of myelination and also regulated by the UPR, decreased with Luman knockdown. In contrast, pro-apoptotic BDNF precursor (proBDNF) levels increased in Luman-deficient SCs at higher ER stress levels, indicating that any protection Luman confers at moderate ER stress levels is lost upon its reduced expression. In conclusion, a connection between Luman and adaptive beneficial ER stress pathways linked to survival and myelination capacity in SCs exists. These Luman-driven cytoprotective mechanisms including survival and myelination open avenues for targeting this pathway in nerve trauma and myelinating disorders.

## INTRODUCTION

Myelin is a complex, multilayered membrane that regulates the speed of electrical impulses traveling along the axon while also providing trophic support to the axon it wraps around (Bouçanova & Chrast, 2020; Bunge, 1994). During myelin production, Schwann cells, glial cells that produce the myelin sheath in the peripheral nervous system (PNS), need to produce high quantities of myelin membrane proteins and cholesterol through the secretory pathway to ensure proper axonal development and neuronal health. Thus, it is not surprising that Schwann cells are highly susceptible to dysfunctions in the secretory pathway, particularly those involved in protein folding and cholesterol biosynthesis (Clayton & Popko, 2016).

The unfolded protein response (UPR) is an in-built cellular mechanism designed to increase protein folding capacity of the endoplasmic reticulum (ER) and maintain lipid homeostasis (Moncan et al., 2021; Schröder & Kaufman, 2005) in response to ER stress initiated by increased volumes of intracellular misfolded protein. UPR initiation involves the binding of misfolded protein to the binding immunoglobulin protein (BiP), releasing them from their silencing roles in association with the three ER-resident master regulatory sensor arms of the UPR. This release of BiP allows activation of the three master regulators: inositol-requiring protein 1α (IRE1α), protein kinase RNA-like ER kinase (PERK), and activating transcription factor 6 (ATF6) (Schröder & Kaufman, 2005; Walter & Ron, 2011). Activated IRE1α catalyzes the unconventional splicing of X-box binding protein mRNA (*Xbp1*) to an active spliced form (*Xbp1s*), upregulating genes involved in ER-associated degradation (ERAD) to increase ER folding capacity (Yoshida et al., 2001). Activated PERK triggers phosphorylation of the α subunit of eukaryotic initiation factor 2 (p-eIF2α), attenuating general protein synthesis to reduce protein load, while selectively translating ATF4 mRNA to initiate gene expression of CCAAT/enhancer-binding protein homologous protein (CHOP), best known as a proapoptotic factor, at chronic stress levels – typically leading to cell death (Z. Liu et al., 2015; Oyadomari & Mori, 2004). Finally, activated ATF6 translocates to the Golgi where cleavage by site-1 and site-2 proteases (S1P and S2P, respectively) releases the amino-terminal cytoplasmic domain of ATF6 which translocates to the nucleus to initiate or alter the expression of genes involved in ER chaperone production and lipid biosynthesis (Haze et al., 1999; Moncan et al., 2021). Activation of any of the canonical UPR pathways stabilizes ER-mediated protein folding mechanisms to improve cellular function. However, constitutive UPR activation resulting from severe or sustained ER stress can activate proapoptotic genes to initiate cell death in an effort to reduce global cellular damage (Urra et al., 2013; Walter & Ron, 2011).

ER stress and activation of the UPR pathways have previously been implicated in the etiology of several PNS demyelinating disorders. Mutations in myelin protein zero (MPZ), for example, can lead to activation of the terminal arm of the UPR by increasing retention of misfolded MPZ in the ER, thereby initiating the PERK pathway (Pennuto et al., 2008; Wrabetz et al., 2006). However, in several cases, the adaptive arm of the UPR system has also been implicated in repairing demyelinated lesions, as well as enhancing the expression of repair-associated markers post-injury. Increasing the levels of spliced *Xbp1* and sustained activation of the IRE1 arm, for example, have been shown to improve motor functions and general clinical outcomes in experimental autoimmune encephalomyelitis mouse model of multiple sclerosis and peripheral nerve injury, respectively (Bracchi-Ricard et al., 2023; Oñate et al., 2016). But the upstream regulators remain largely unknown.

We have previously revealed that the axonal ER transmembrane-associated transcription factor Luman/CREB3, herein called Luman, functions as a critical sensor and retrograde regulator of axon repair in injured axons via its regulation of neuronal ER stress, demonstrating a connection between axon regeneration and modulation of the UPR (Ying et al., 2014, 2015). The Luman-dependent introduction of low-level beneficial ER stress that leads to an adaptive induction of the UPR and cholesterol biosynthesis during peripheral nerve injury, has the capacity to promote axon regeneration in sensory neurons (Ying et al., 2015). Luman is implicated in ER stress response and the UPR through its sequence and activation similarity with ATF6 (R. Lu et al., 1997) and its capacity to bind ER stress response elements in target genes (DenBoer et al., 2005). Unlike ATF6 though, Luman is not proteolytically cleaved/activated, nor its expression levels altered, in response to the UPR inducers tunicamycin or thapsigargin (DenBoer et al., 2005; Ying et al., 2015). This suggests that the pathway involved in Luman-mediated UPR activation varies from the currently known ER stress responses and provides a tool to induce the UPR without altering the Luman activation state. Additional physiological roles for Luman include mediating Lkn-1 cytokine-induced migration of monocytes (Jang et al., 2007), control of folliculogenesis and synthesis of progesterone and estradiol (Zhao et al., 2016), and suppression of tumor progression (He et al., 2024).

Despite the various reported functions of Luman in a variety of cell types, little is known about the role Luman plays in Schwann cells. The combination of previous research we have conducted revealing a role for Luman in sensory neuron axon regeneration through the injury-induced UPR and cholesterol biosynthesis pathways, the latter a critical component of myelin (Saher & Simons, 2010), and aforementioned studies describing the role ER stress plays in demyelinating disorders, myelin synthesis, and myelinating glial cell survival and maturation, led us to posit that Luman is also be a key regulator of the myelination capacity and viability of Schwann cells. Characterization of Luman’s role in these areas, especially under varying levels of ER stress, would help us gain a better understanding about how Luman regulates the adaptive UPR mechanisms that could be targeted to enhance the intrinsic cellular mechanisms involved in neural repair.

This study tested the hypothesis that myelination of sensory axons is regulated by Luman-associated control of the myelination capacity and viability of Schwann cells. We demonstrate that Luman positively regulates these parameters in the context of low-level beneficial stress brought about by activation of downstream UPR-associated pathways, including brain-derived neurotrophic factor (BDNF) and sterol expression.

## MATERIALS AND METHODS

### Animals

Male Wistar rats (2-3 months of age) were used for primary Schwann cell cultures and P3 Wistar rat neonates were used for isolating DRGs for co-cultures. All animals were obtained from Charles River Laboratories (Senneville, QC, Canada). All animal procedures conducted were approved by the University of Saskatchewan Animal Research Ethics Board and adhered to Canadian Council on Animal Care guidelines.

### Primary Schwann cell cultures

Primary Schwann cells were isolated from the sciatic nerves of adult male Wistar rats as previously described (Kaewkhaw et al., 2012) with some modifications. Briefly, rat sciatic nerves were excised surgically, maintained in Leibovitz’s L-15 medium (Invitrogen), and the tissues were treated with 1 mg mL^-1^ Type I collagenase (Invitrogen) and 2.5% trypsin in Puck’s solution (Sigma-Aldrich). Culture vessels were pre-treated with 25 µg mL^-1^ poly-L-lysine (Sigma-Aldrich) and 1 µg mL^-1^ laminin (Sigma-Aldrich). Schwann cells were cultured in Dulbecco’s modified eagle medium with D-valine substituted for L-valine (Boca Scientific, Dedham, Massachusetts, USA) supplemented with 10% fetal bovine serum (FBS; Invitrogen), 100 U mL^-1^ penicillin/100 µg mL^-1^ streptomycin and 0.25 µg mL^-1^ amphotericin B (Sigma-Aldrich), 2mM L-glutamine (Invitrogen), 1% (v/v) N2 supplement (Invitrogen), 10 µg mL^-1^ bovine pituitary extract (Invitrogen), 5 mM forskolin (Sigma-Aldrich), and 10 ng mL^-1^ neuregulin/NRG1-β1 (R&D Systems) in a humidified 5% CO_2_ incubator at 37°C. Cultured primary Schwann cells were passaged no more than three times before being used.

### P3 DRG/SC co-cultures

DRG explants were isolated from P3 Wistar rat neonates and cultured individually on glass coverslips treated with 25 µg mL^-1^ poly-L-lysine, 10 µg mL^-1^ laminin, Type I collagen (Sigma-Aldrich), and Matrigel (Corning) in Neurobasal-A medium (Invitrogen) supplemented with 20% FBS, 100 U mL^-1^ penicillin/100 µg mL^-1^ streptomycin and 0.25 µg mL^-1^ amphotericin B, 1% (v/v) B-27 supplement (Invitrogen), 50 ng mL^-1^ nerve growth factor 2.5S (NGF; Cedarlane, Burlington, ON, Canada), and 7 µM cytosine arabinoside (Ara-C; Sigma-Aldrich). Culture media was changed every 2 days and Ara-C was removed from the medium at 6 days *in vitro* (DIV). At DIV 8, Schwann cells were co-cultured with the DRG explants in 1:1 DMEM with Ham’s F-12 (Invitrogen) supplemented with 10% FBS, 100 U mL^-1^ penicillin/100 µg mL^-1^ streptomycin and 0.25 µg mL^-1^ amphotericin B, 1% (v/v) N2 supplement, 50 ng mL^-1^ NGF, 5 mM forskolin, and 10 ng mL^-1^ neuregulin/NRG1-β1. Once the Schwann cells aligned with the DRG neurites, media was changed to 1:1 DMEM with Ham’s F-12 supplemented with 5% FBS, 100 U mL^-1^ penicillin/100 µg mL^-1^ streptomycin and 0.25 µg mL^-1^ amphotericin B, 1% (v/v) N2 supplement, 50 ng mL^-1^ NGF, and 0.1% (w/v) L-ascorbic acid (Sigma-Aldrich) to initiate myelination. Cultures were maintained for 21 days after initiation of myelination.

### siRNA transfection

Knockdown of Luman expression was achieved through siRNA transfection. Poly-L-lysine- and laminin-coated 96-well plates were seeded with Schwann cells at a density of 10^4^ cells/well 48 hours prior to transfection. Cultures were transfected using the TriFECTa RNAi Kit (Integrated DNA Technologies, Coralville, IA, USA) per manufacturer’s recommended protocol. DsiRNA sequences used included rn.Ri.Creb3.13.1, rn.Ri.Creb3.13.2, rn.Ri.Creb3.13.3; negative control (scrambled sequence control - Integrated DNA Technologies). DsiRNA were transfected using Lipofectamine RNAiMAX Transfection Reagent (Life Technologies). Transfection efficiency of siRNA constructs was assessed by transfection with a TYE 563-conjugated transfection control (10 nM) followed by quantification of cellular uptake through fluorescence imaging using the Zeiss Axio Observer with Axiocam 305 fluorescence microscope at 4 hours, 24 hours, 48 hours, 72 hours, and 96 hours post-transfection. Luman knockdown efficiency following siRNA transfection was assessed by quantitative real time PCR (RT-qPCR) and western blot 48 hours post-transfection.

### Adenoviral transduction

Overexpression of Luman was done by transduction of cells with adenovirus containing the rat Luman open reading frame prepared using the Adeno-X Viral DNA Vector (Clontech, Mountain View, CA, USA) and purified using the Adeno-X Expression System (Clontech) as per Ying, et al. (2014). For *in vitro* transductions, Schwann cells were infected 24 hours prior to co-culture initiation with 3.2×10^8^ pfu/µL of either Luman AdV or lac Z AdV (negative control AdV expressing *Escherichia coli* β-galactosidase) at a multiplicity of infection (MOI) of 75. An untreated culture setup with no AdV transduced was also maintained. Transduction efficiency of adenoviruses 24 hours and 48 hours post-transduction was assessed by fluorescent detection in transduction controls of GFP and RFP tags conjugated to the N-terminus and C-terminus of the Luman protein, respectively. Luman overexpression efficiency following AdV transduction was assessed by RT-qPCR and western blot 24 hours post-transduction.

### Tunicamycin treatment

Tunicamycin (Sigma-Aldrich) treatment was done to initiate ER stress on cultures and treatment was performed concurrent to siRNA transfection. Tunicamycin was resuspended in complete growth medium and used at final concentrations of 2, 10, and 100 ng mL^-1^.

### Cell viability assay and apoptosis assay

Cell viability was assessed in cultures using the PrestoBlue Cell Viability Reagent (Invitrogen, A13261) per manufacturer’s recommended protocol. Briefly, the reagent was added at a 1:10 dilution to each well of a 96-well plate with Schwann cell cultures and incubated for 30 minutes in a CO_2_ incubator at 37°C. Plates were read using a SpectraMax M2 Multimode Microplate Reader (Molecular Devices, San Jose, CA, USA) by detecting fluorescence levels at an excitation wavelength of 544 nm and emission wavelength of 590 nm, with a cutoff at 590 nm. Background fluorescence was subtracted from no-cell controls. Percent viability was calculated by dividing mean absorbance of sample over mean absorbance of blank multiplied by 100. Apoptotic cell death was evaluated using the Apoptosis/Necrosis Assay Kit (Abcam, ab176749) per manufacturer’s recommended protocol. Briefly, stock components were diluted to 1X in culture medium, added to the wells of 96-well plates with Schwann cell cultures, and incubated for 1 hour in a CO_2_ incubator at 37°C. Apoptotic cells were stained with Apopxin Green and detected using the FITC filter and live cells were stained with Cytocalcein Violet 450 and detected using the DAPI channel of the Zeiss Axio Observer fluorescence microscope. Images were analyzed using FIJI Image J (Schindelin et al., 2012) and the level of apoptosis was quantified by calculating the percentage of apoptotic cells over total number of cells. Ten areas were analyzed per data group.

### Cholesterol assay

The amount of total cholesterol produced by the cultures was measured 48 hours post-transfection using the Amplex Red Cholesterol Assay Kit (Invitrogen, A12216) per manufacturer’s recommended protocol. Data were normalized to protein content, with total cholesterol levels represented as µg of total cholesterol per mg of protein quantified using Bradford assay (Bio-Rad).

### Quantitative real-time polymerase chain reaction (RT-qPCR)

Total RNA was extracted from cell cultures using the RNeasy Plus Mini Kit (Qiagen, 74134) following manufacturer’s protocol. RT-qPCR was conducted using the QuantiNova SYBR Green RT-PCR Kit (Qiagen, 208354) on a QuantStudio 3 Real-Time PCR System (Applied Biosystems, Waltham, MA, USA). Thermocycling conditions were 50°C for 10 minutes, 95°C for minutes, followed by 40 cycles of 95°C for 5 seconds and 60°C for 10 seconds. Relative expression levels of specific genes were calculated using the 2^-ΔΔCt^ method with 18S rRNA serving as an internal control. The primer pairs used were: *Luman* (forward) 5’- TGTGCCCGCTGAGTATGTTG-3’ (reverse) 5’-AGAAGGTCGGAGCCTGAGAA-3’; *Mbp* (forward) 5’-TCCGGAGGTTCAGGTGCACG-3’ (reverse) 5’-CTGGACTCTCACAGCTGCCC- 3’; *Xbp1* (forward) 5’-CAGACTACGTGCACCTCTGC-3’ (reverse) 5’- CAGGGTCCAACTTGTCCAGAAT-3’; *Xbp1s* (forward) 5’-GCTGAGTCCGCAGCAGGT-3’ (reverse) 5’-CAGGGTCCAACTTGTCCAGAAT-3’; *Chop* (forward) 5’- TGGAAGCCTGGTATGAGGAC-3’ (reverse) 5’-TGCCACTTTCCTCTCGTTCT-3’; *Srebf1* (forward) 5’-CGCTACCGTTCCTCTATCA-3’ (reverse) 5’-CTCCTCCACTGCCACAAG-3’; *Bdnf* (forward) 5’-GCCTCCTCTGCTCTTTCT-3’ (reverse) 5’-GCCGTTACCCACTCACTA-3’; *18S rRNA* (forward) 5’-TCCTTTGGTCGCTCGCTCCT-3’ (reverse) 5’-TGCTGCCTTCCTTGGATGTG-3’. RT-qPCR analysis was performed in triplicate and data were reported as fold change levels.

### Western blot analysis

Total protein was extracted from cultures using CelLytic M Lysis Buffer (Sigma-Aldrich) supplemented with Protease and Phosphatase Inhibitor Cocktail (Sigma-Aldrich) according to manufacturer’s recommended protocol and quantified using a Nanodrop 2000 Spectophotometer (Thermo Scientific). For western blotting, 10 µg of total protein per sample was separated using SDS-PAGE on 10% stain-free polyacrylamide gels (Bio-Rad) and then transferred to a PVDF membrane (Bio-Rad) prior to blocking for one hour at room temperature with either 10% SeaBlock Blocking Buffer (Thermo Scientific) in 1X PBS with 0.1% Tween-20 (PBST) for BDNF immunoblots, 5% goat serum in 1X TBS with 0.1% Tween-20 (TBST) for pIRE1α immunoblots, or 5% goat serum in 1X PBST for the rest of the immunoblots. The immunoblots were then incubated in primary antibodies overnight at 4°C in their respective blocking solutions. Primary antibodies used include rabbit anti-Luman (1:800; Misra Laboratory, University of Saskatchewan; Hasmatali et al., 2019), rabbit anti-MBP (1:1000; Abcam), mouse anti-NF200 (1:1000; Sigma-Aldrich), rabbit anti-BDNF (1:1000; Abcam), rabbit anti-CHOP (1:1000; Santa Cruz Biotechnology, Inc., Dallas, TX, USA), and rabbit anti-p-IRE1α (1:1000; Abcam). Immunoblots were then incubated with horseradish peroxidase-conjugated secondary antibodies (1:3000; Bio-Rad) for one hour at room temperature and western blots were visualized with Bio-Rad Clarity Western enhanced chemiluminescence reagents using the Bio-Rad ChemiDoc imaging system. To quantify relative differences in protein expression among different treatment setups, bands in the immunoblots were subjected to densitometric analysis with the Bio-Rad ImageLab 6.0.1 software using total protein levels as internal loading controls.

### Immunostaining

Cell cultures were prepared and fixed in 2% paraformaldehyde in PBS for 30 minutes at room temperature and stored in PBS overnight at 4°C. After fixing, cultures were blocked with SeaBlock blocking buffer for one hour. Cultures were then incubated with primary antibodies overnight at 4°C, and then incubated with fluorescent-dye-conjugated secondary antibodies for one hour at room temperature. Primary antibodies used include rabbit anti-Luman antibody (1:400; Misra Laboratory, University of Saskatchewan; Hasmatali et al., 2019), rabbit anti-MBP antibody (1:250, Abcam), and mouse anti-neurofilament 200 (1:400, Sigma-Aldrich). Secondary antibodies used include goat anti-mouse IgG conjugated to AlexaFluor488 (1:750; Abcam) and donkey anti-rabbit IgG conjugated to Cy3 (1:3000; Jackson ImmunoResearch Labs, West Grove, PA, USA). Nuclear staining was done with DAPI (1:5000; Life Technologies). Images were obtained using the Zeiss Axio Imager M.1 fluorescence microscope and analyzed and further processed using the Zen 3.4 Blue Edition software (Carl Zeiss, ON, Canada) and FIJI ImageJ.

### Analysis and quantification of immunofluorescence signal

To quantify the total area covered by MBP, the NF200-positive zones were used to generate a mask to define the mature axonal regions. The area covered by MBP was then determined and the MBP/NF200 ratios were calculated to determine the amount of MBP expressed over a number of mature neurites. The values were then normalized against the mean value of the average ratios for the untreated condition.

To ensure the relative changes in immunofluorescence signal among experimental groups were analyzed accurately, coverslips from untreated, negative control, and experimental groups were processed alongside each other under identical conditions. Images were acquired under identical exposure levels, blinded using the Blind Analysis Tools plugin in FIJI ImageJ, and downstream analysis was performed in parallel. All analysis was done was done in a manner blinded to the condition.

### Statistical analysis

Data are expressed as mean ± SD for the indicated number of observations. One-way ANOVA with post-hoc Tukey’s test for multiple comparisons was utilized to assess data from cell viability, expression level comparisons, and myelination assay studies to compare untreated and negative control groups against treatments. Two-way ANOVA with post-hoc Tukey’s test for multiple comparisons was used to assess data from ER stress-induced assays to compare effects of different levels of ER stress and presence of siRNA in specific setups. Statistical significance was accepted at p <0.05. Statistical analysis was performed using GraphPad Prism 9.0.0 software (San Diego, CA, USA).

## RESULTS

### Luman’s impact on Schwann cell viability and initiation of ER stress-associated apoptotic events

To determine the effect of altered Luman expression on Schwann cells, an siRNA pool with three siRNA constructs was designed to target various regions in the *Luman* mRNA sequence. Transfection efficiency in primary cultures of Schwann cells derived from adult rat sciatic nerves was evaluated using a positive control siRNA with a TYE 563 fluorescent label. Peak transfection was observed 48 hours post-transfection (Figure S1A, B). Quantification of *Luman* transcript expression (Figure S1C) and Luman protein levels (Figure S1D, E) showed a significant drop by 95% and 70% respectively by the peak transfection timepoint relative to levels in untransfected Schwann cells or cells transfected with control, nontargeting siRNA, validating the transfection protocol. The percent viability of Schwann cells following transfection with Luman siRNA was significantly reduced, indicating the importance of Luman expression in maintaining Schwann cell survival *in vitro* (Figure 1A). No significant difference in Schwann cell percent viability was observed between untransfected and control siRNA-transfected groups.

**Figure 1.**
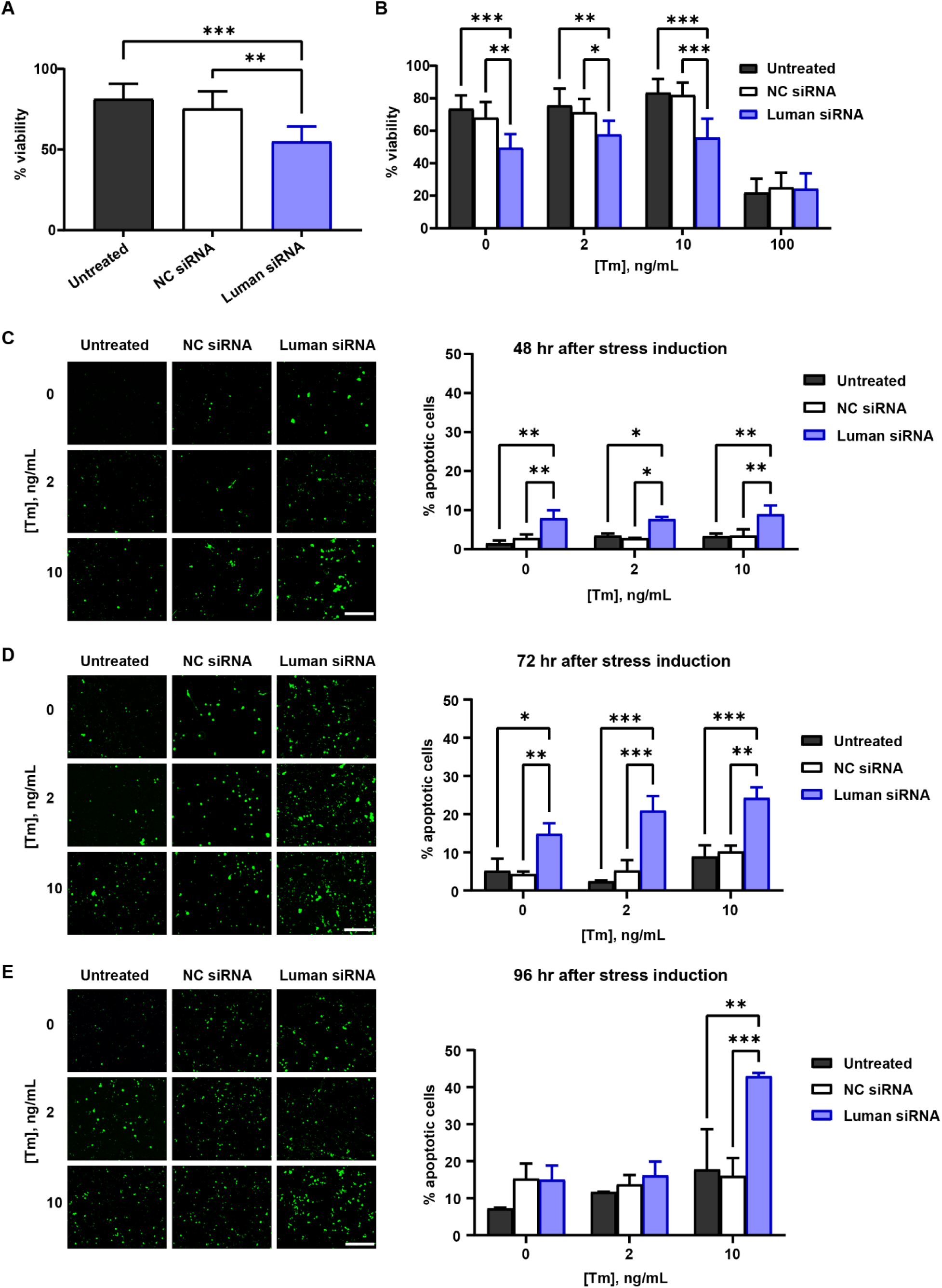
Luman maintains Schwann cell viability and prevents initiation of ER stress-induced apoptotic events. Percentage of viable Schwann cells determined by PrestoBlue assay 48 hours after transfection with either no siRNA (untreated), negative control non-targeting siRNA (NC siRNA), or 3 Luman siRNA constructs (Luman siRNA) without (A) and with (B) ER stress induction by addition of 0, 2, 10, or 100 ng mL^-1^ of tunicamycin (Tm). Schwann cells viability 48 hours (C), 72 hours (D), and 96 hours (E) after siRNA transfection with or without ER stress induction by addition of 0, 2, or 10 ng mL^-1^ of tunicamycin. Percentage of apoptotic cells is number of apoptotic cells (green) over total number of cells in each plate. Ten areas analyzed per data group. Images are representative. Scale bar = 100 µm. All data derived from *N* = 6 (A) or *N = 3* (B-E) experimental repeats for each condition. Data expressed as mean ± SD. One-way (A) or two-way ANOVA (B-E) with post-hoc Tukey’s test (*p<0.05, **p<0.01, ***p<0.001).

Ying et al. (2015) previously revealed that Luman exerted its impacts on sensory neuron axon regeneration through regulation of the unfolded protein response in response to injury-initiated ER stress. Furthermore, reduced axon outgrowth in injured sensory neurons in response to Luman siRNA transfection was able to be rescued by ER stress induction with low levels of tunicamycin, an N-linked glycosylation inhibitor that can cause G1 cell cycle arrest (C. Han et al., 2013). Importantly, we demonstrate that tunicamycin does not alter Luman expression in Schwann cells (Figure S2), which has also been observed in different cell types in previous studies (DenBoer et al., 2005; Ying et al., 2015), allowing for its combinatorial use in our study.

In order to assess the effect of altering Luman expression on Schwann cell viability in the context of increasing levels of ER stress, ER stress was induced by supplementation of the culture medium with various concentrations of tunicamycin. Schwann cell viability following siRNA transfection under unstressed (0 ng·mL^-1^ tunicamycin), mild (2 ng·mL^-1^ tunicamycin), moderate (10 ng·mL^-1^ tunicamycin), and severe (100 ng·mL^-1^ tunicamycin) ER stress levels was evaluated. It should be noted, that different cell types have different levels of sensitivity to ER stress and that the terms “mild”, “moderate”, and “severe” only serve as nominal metrics of relative ER stress intensity for Schwann cells rather than a measure of absolute intensity. Mild and moderate ER stress conditions were an attempt to replicate *in vivo* activation of the unfolded protein response at concentrations that were low enough to initiate the adaptive UPR arm in a manner akin to its induction in sensory neurons in response to peripheral nerve injury and without tunicamycin itself inducing neuronal death, but high enough to have a cellular effect linked to successful axon regeneration (Ying et al., 2014, 2015).

Luman knockdown reduced Schwann cell survival under mild and moderate ER stress levels, relative to the transfection controls by an average of 23.7% and 33.1% respectively, while severe ER stress levels significantly reduced Schwann cell survival to 22% independent of Luman expression levels (Figure 1B). As a result, further assays involving ER stress induction solely involved the mild and moderate ER stress levels induced by tunicamycin, so as to distinguish Luman’s role.

We next explored the nature of Schwann cell death following Luman knockdown. There was a significant increase in the percentage of apoptotic cells both 48 hours and 72 hours following Luman knockdown in Schwann cells (Figure 1C, D). This was observable under unstressed conditions, as well as following mild and moderate ER stress induction. Although this is the case, 96 hours post-siRNA transfection, the percentage of apoptotic cells was only increased in Schwann cells where Luman was knocked down in moderately stressed cells (Figure 1E). There was no significant increase in rate of apoptosis under the lower ER stress conditions when Luman is knocked down. There was also no detectable secondary apoptosis or necrotic cell death within the set timeframes of the assay (data not shown).

### Regulation of Schwann cell-mediated myelination of sensory axons *in vitro* by Luman

Myelination assays involving Schwann cells and dorsal root ganglion (DRG) neuron co-cultures are a well-established *in vitro* system to observe the impact of select molecules, such as Luman, on myelination. However, to observe myelination, it necessitates maintaining the co-cultures for up to 21-24 days after establishment (Taveggia & Bolino, 2018). Due to our finding that reducing Luman expression with siRNA resulted in increased cell death and reduced Schwann cell numbers shortly after transfection, this approach could not be repurposed to examine the effects of reduced Luman expression on myelinating capacity of Schwann cells on sensory neurons *in vitro*. As a result, an adenoviral vector-based method designed to overexpress Luman was used on Schwann cells just prior to establishing the myelination co-cultures with DRG neurons. Adenoviral vectors have been demonstrated to successfully overexpress genes in Schwann cells without major cytopathic effects for up to 45 days (Shy et al., 1995). To visually track the localization of the Luman construct in Schwann cells after transduction, a construct previously designed and used in our lab was employed. This construct has a GFP tag fused to the N-terminus of the Luman protein, which translocates after being cleaved in the Golgi apparatus into the nucleus, and an RFP tag was fused to the C-terminus of the Luman protein, the latter remaining in the ER after activation (Ying et al., 2014). Both tags were visible intracellularly by 48 hours post-transduction and their expression levels correlated with intracellular Luman levels after transduction (Figure S3A). Accordingly, *Luman* transcript (Figure S3B) and Luman protein levels (Figure S3C, D) were found to increase in Luman AdV-transduced Schwann cells, including single-tagged and non-tagged Luman adenoviral constructs, thus validating the overexpression protocol.

To examine whether increased Luman expression in Schwann cells enhances myelination, Schwann cells were transduced with either no adenovirus (vehicle), control non-tagged adenovirus (expressing lacZ), or Luman adenovirus (Luman AdV) 24 hours prior to establishing the co-cultures with DRGs. Once the Schwann cells aligned to the DRG neurites, we added L-ascorbic acid to the culture medium. Both alignment and L-ascorbic acid are necessary to induce peripheral nerve myelination *in vitro* (Eldridge et al., 1987; Kleitman et al., 1998). The cultures were then maintained for 21 days before staining for myelin basic protein (MBP) and neurofilament-200 (NF200) to evaluate myelination outcomes.

DRG/Schwann cell co-cultures containing Schwann cells transduced with Luman AdV exhibited a significant increase in MBP expression, an indicator of increased compact myelin by Schwann cells (Figure 2A) (Weil et al., 2016). Specifically, myelination of mature NF200-positive neurofilaments, as evidenced by the colocalization of MBP and NF200, increased in co-cultures where the Schwann cells were transduced with Luman AdV (Figure 2B) when assessed 21 days post initiation of myelination. Levels of *Mbp* transcript also increased (Figure 2C) in myelinating Schwann cells following Luman overexpression. MBP levels were shown to increase with respect to NF200 levels, which remained constant (Figure 2D, E). This indicates that overexpression of Luman in Schwann cells increased their capacity to myelinate mature axons but did not appear to impact neurite levels in the cultures.

**Figure 2.**
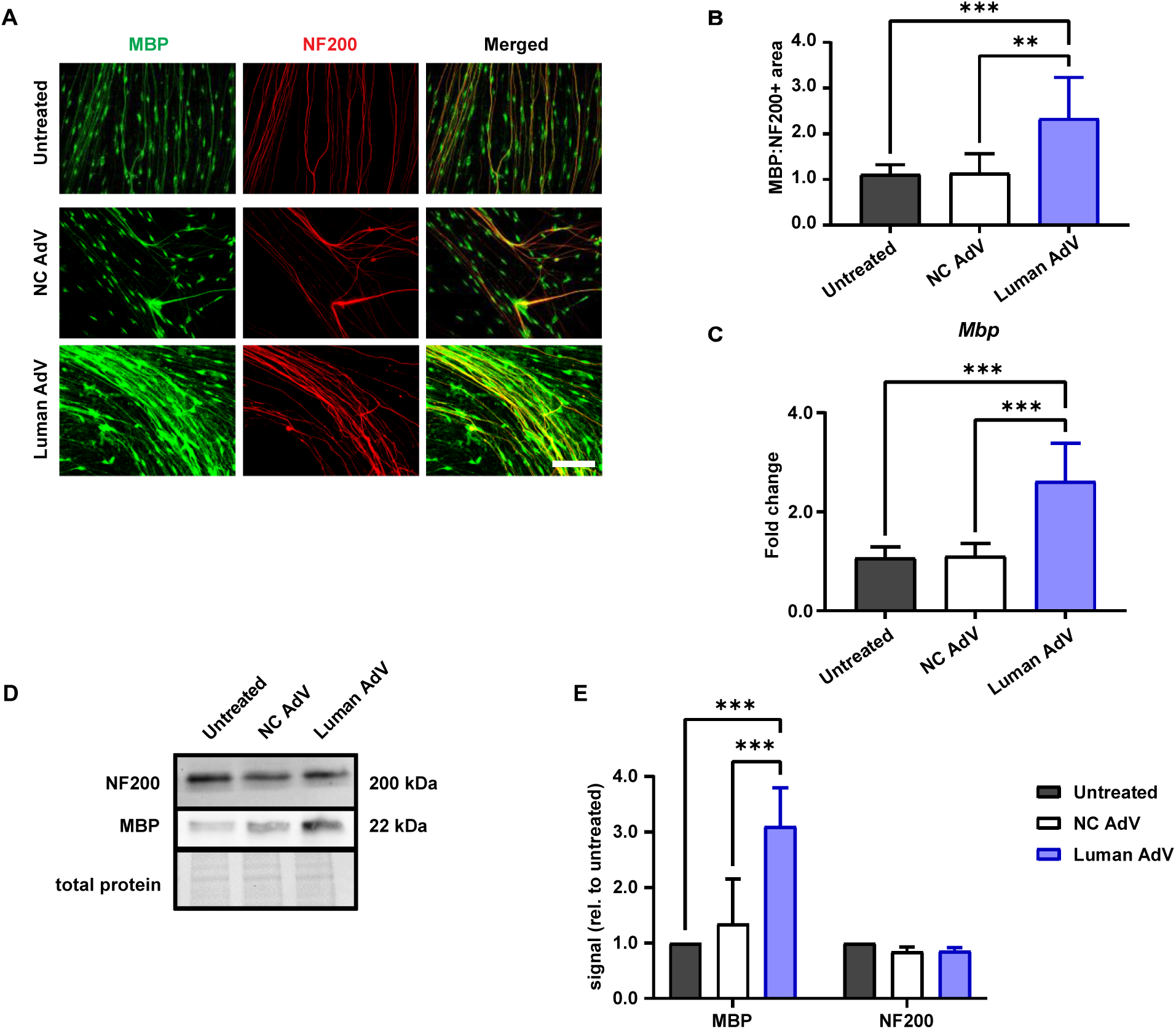
Luman positively regulates Schwann cell-mediated myelination of sensory neurons *in vitro*. (A) Representative images of sensory neuron/Schwann cell myelination co-cultures 21 days after initiation of myelination. Schwann cells were transduced with no adenovirus (untreated), lacZ negative control adenovirus (NC AdV), or Luman-overexpressing adenovirus (Luman AdV) 24 hours prior to initiation of co-cultures. Cultures were dually stained for myelin basic protein (MBP) (green) to visualize compact myelin structures and neurofilament-200 (NF200) (red) to visualize mature sensory neuron axons. Scale bar = 20 µm. (B) Ratios of MBP immunofluorescence area relative to NF200 immunofluorescence as normalized to mean untreated groups ratio. (C) RT-qPCR quantification of fold change of *Mbp* transcript levels in experimental groups (as indicated) from myelination co-cultures 21 days post-myelination initiation. (D) Representative immunoblots of protein isolated from myelination co-cultures with DRG explants removed just prior to lysis as in (C) and probed for MBP (22 kDa) and NF200 (200 kDa). Densitometric analysis of band intensities was performed and data were normalized to total protein levels in each lane. (E) Band intensities from MBP-and NF200-probed immunoblots normalized to the mean band intensities in untreated groups. All data derived from *N* = 3 experimental repeats/condition. Data expressed as mean ± SD. One-way ANOVA with post-hoc Tukey’s test (*p<0.05, **p<0.01, ***p<0.001).

### Impact of Luman on the IRE1 and PERK arms of the UPR in Schwann cells

Luman’s ability to regulate axon regeneration has been linked to its regulation of the UPR pathway (Ying et al., 2014, 2015). To examine the role of the ER stress-associated transcription factor Luman in Schwann cells, we knocked down Luman expression using siRNA-mediated transfection as above and observed the differences in transcript and protein levels of various IRE1 and PERK pathway-associated markers within and outside the context of ER stress induced by tunicamycin 48 hours previously.

We observed a significant increase in *Xbp1* mRNA levels in Schwann cells following knockdown of Luman expression under unstressed conditions (Figure 3A. In contrast, following mild and moderate ER stress induction, Luman knockdown led to a significant decrease in *Xbp1* mRNA levels compared to untransfected and control siRNA-transfected groups with levels heightened in these control groups in response to mild stress (Figure 3A). The levels of spliced *Xbp1* transcript (*Xbp1s*) follow a similar trend to changes in *Xbp1* transcript levels; following ER stress induction and Luman knockdown, unstressed Schwann cells also exhibit an increased level of *Xbp1s*. But following mild and moderate ER stress induction, *Xbp1s* levels drop in the Luman-knocked down Schwann cells compared to untransfected and control siRNA-transfected groups (Figure 3B). This contrasts with the marked increase of *Xbp1s* transcript levels in moderately ER-stressed Schwann cells, which, although being significantly elevated in all treatment groups relative to unstressed or mildly stress cells, its expression is still significantly reduced in response to Luman siRNA treatment (Figure 3B).

**Figure 3.**
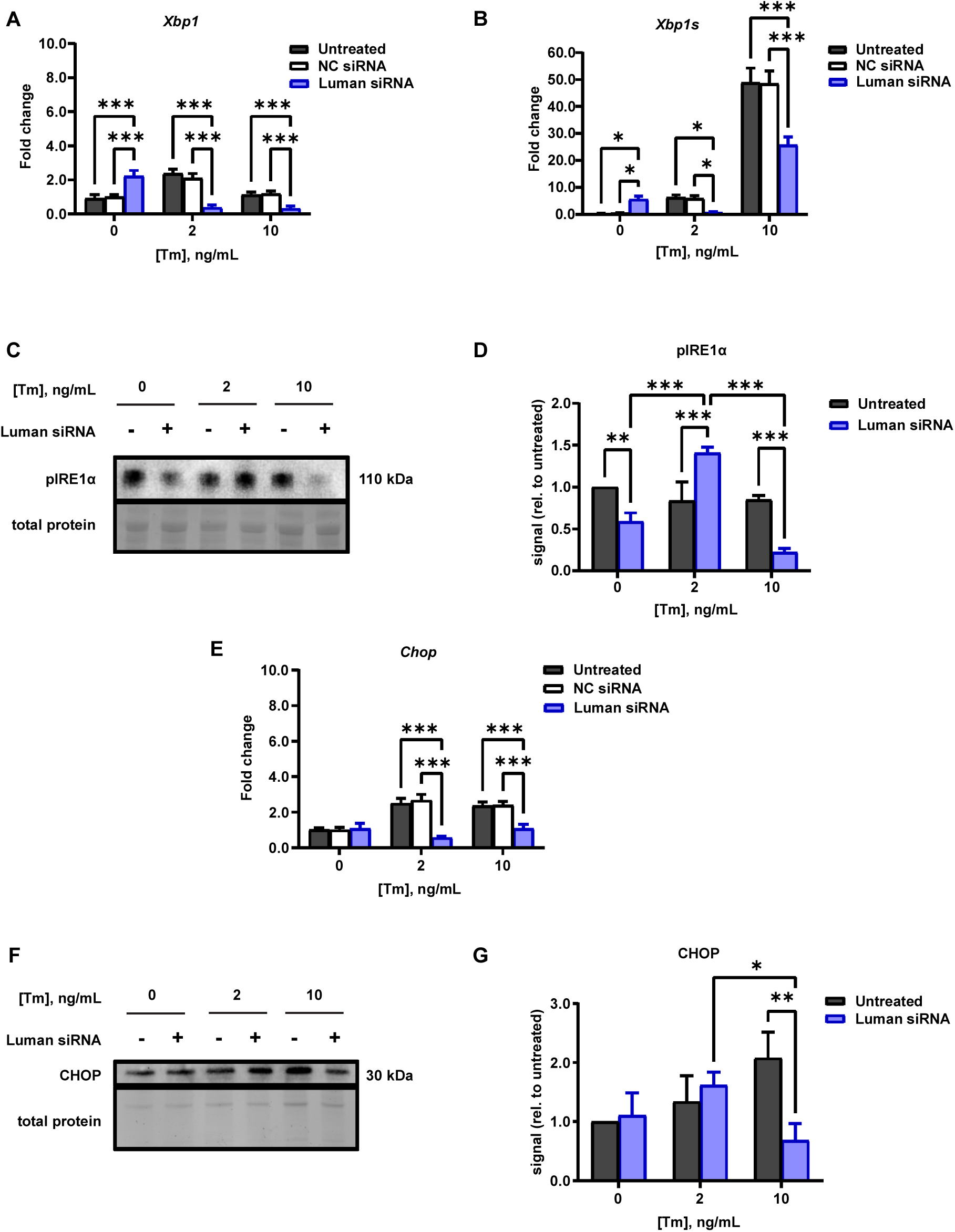
Luman regulates the IRE1 and PERK arms of the unfolded protein response in Schwann cells. RT-qPCR quantification of fold change of *Xbp1* (A), spliced *Xbp1* [*Xbp1s*] (B), and *Chop* (E) transcript levels in Schwann cells 48 hours after siRNA transfection with or without ER stress induction by addition of 0, 2, or 10 ng·mL^-1^ of tunicamycin. Representative immunoblots of protein isolated from Schwann cells 48 hours after siRNA transfection with or without ER stress induction by addition of 0, 2, or 10 ng·mL^-1^ of tunicamycin and probed for pIRE1α (C) at 110 kDa and CHOP (F) at 30 kDa. Densitometric analysis of band intensities was performed and data were normalized to total protein levels in each lane. Band intensities from pIRE1α- (D) or CHOP- (G) probed immunoblots were then normalized to the mean band intensities in untreated groups. All data derived from *N* = 3 experimental repeats/condition. Data expressed as mean ± SD. Two-way ANOVA with post-hoc Tukey’s test (*p<0.05, **p<0.01, ***p<0.001).

In order to evaluate UPR pathway protein level variations following Luman knockdown and ER stress induction in Schwann cells, the levels of phosphorylated IRE1α protein were quantified under various conditions. Levels of pIRE1α decreased following Luman knockdown in both unstressed and moderately stressed Schwann cells. However, pIRE1α protein levels increased in mildly stressed Schwann cells following Luman knockdown compared to the untreated group suggesting that Luman serves to dampen this response in mildly stressed cells (Figure 3C, D). IRE1 activation is generally associated with higher expression of pIRE1α and subsequent downstream IRE1 products, including XBP1, XBP1s, and BDNF, among others (Saito et al., 2018). While expression level differences between pIRE1α and XBP1/XBP1s would be predicted, due to the upstream nature of the IRE1α protein, it would be natural to assume activation of IRE1 via phosphorylation would also positively correlate with XBP1/XBP1s levels. Finally, there was a subsequent decrease of pIRE1α in Luman-deficient Schwann cells under moderate ER stress conditions, which when taken together with the similarly decreased levels of *Xbp1* and *Xbp1s* transcripts under these conditions, supports that there is a role for Luman in the regulation of the IRE1 arm of the UPR/ER stress response as the cells adapt to the stress.

The PERK arm of the pathway, when persistently activated, has been linked to promotion of ER stress-induced cellular death either through activation of the CHOP-mediated pathway or suppression of the cellular inhibitors of apoptosis (cIAP) protein machinery (Hamanaka et al., 2009; Hiramatsu et al., 2014). Given the role of Luman in Schwann cell survival described above, the influence of Luman expression on PERK activity in Schwann cells was evaluated by quantitation of both *Chop* transcript and CHOP protein levels following Luman knockdown and/or ER stress induction by tunicamycin. We observed a significant decrease in *Chop* transcript levels in both mildly and moderately stressed Schwann cells following Luman expression knockdown compared to cells with normal baseline Luman levels and did not observe any significant influence of Luman expression level changes on *Chop* in unstressed cells (Figure 3E). Similarly, at the protein level, we observed no significant change in CHOP expression levels following Luman knockdown under unstressed conditions. Luman knockdown only significantly decreased CHOP levels under moderate ER stress conditions compared to untreated groups (Figure 3F, G), suggesting its regulation of this pathway at the protein level may only emerge at moderate to higher levels of ER stress.

### Impact of Luman on total cholesterol production at varying levels of ER stress in Schwann cells

We have previously shown that Luman knockdown significantly decreases the expression of cholesterol production-associated genes (such as *Insig1* and SREBP), as well as the total and free cholesterol production in injury-conditioned UPR-activated sensory neurons (Ying, et al., 2015). Since Schwann cells rely heavily on *de novo* synthesis of cholesterol for the production of compact myelin membranes (Montani et al., 2018; Saher et al., 2009), we aimed to determine how Luman regulates cholesterol synthesis and how it affects overall cholesterol production in Schwann cells within and outside the context of the unfolded protein response. Luman knockdown decreased the expression of the major sterol regulatory element binding transcription factor *Srebf1* across all experimental ER stress conditions used (Figure 4A). In addition to this, the total cholesterol production in Schwann cells is also reduced following Luman knockdown (Figure 4B). This general pattern can also be observed following mild and moderate ER stress levels (Figure 4C). However, at severe ER stress levels (100 ng·mL^-1^ tunicamycin), Luman expression ceases to significantly alter total cholesterol levels as they reach the same levels detected in siRNA negative control groups (Figure 4C).

**Figure 4.**
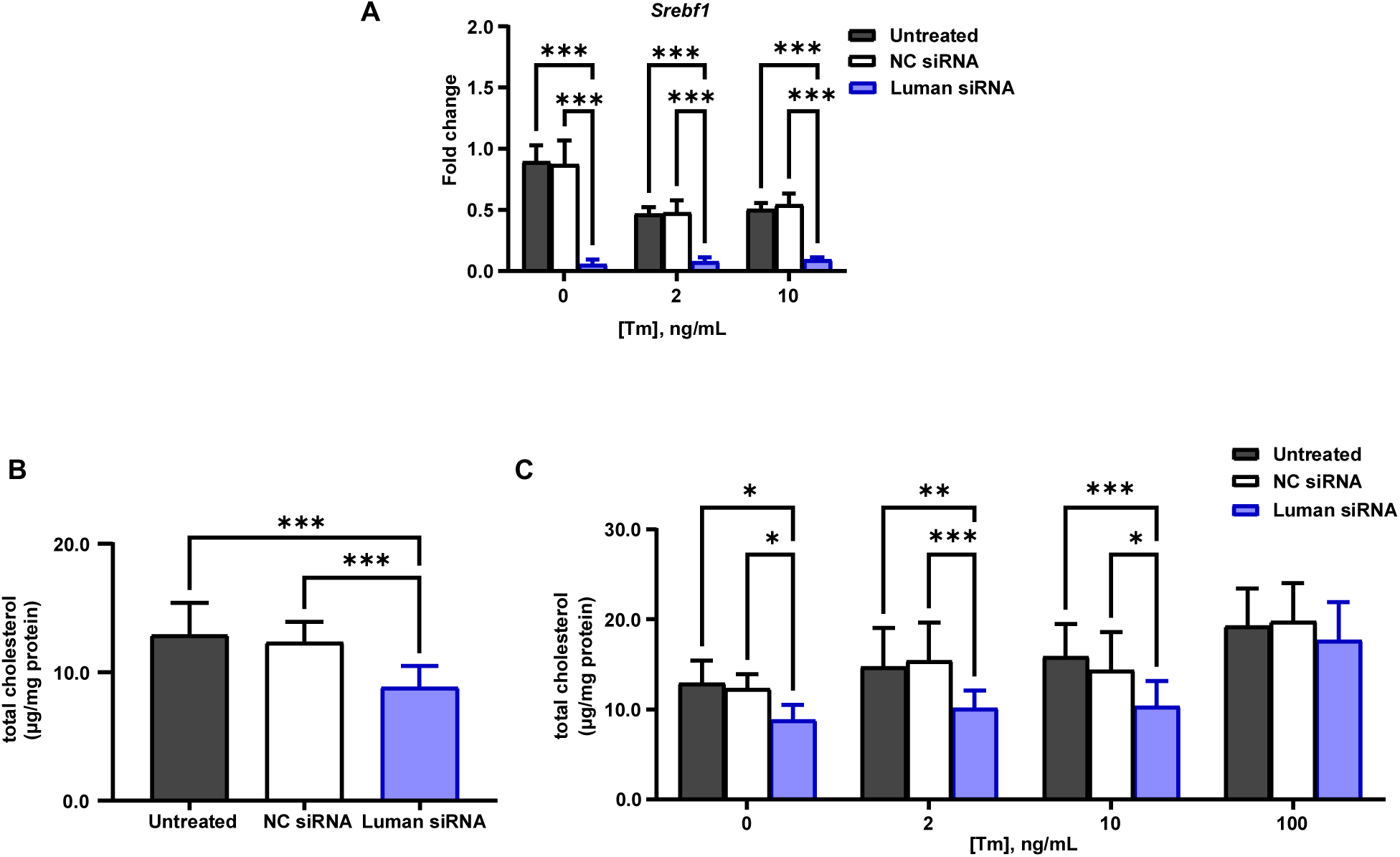
Luman expression positively regulates total cholesterol production at varying levels of ER stress in Schwann cells. (A) Quantification of fold change using RT-qPCR of *Srebf1* transcript levels in Schwann cells 48 hours after siRNA transfection with or without ER stress induction by addition of 0, 2, or 10 ng·mL^-1^ of tunicamycin. Total cholesterol quantification using the Amplex Red Cholesterol Assay Kit (Invitrogen). Fluorescence was measured (545 nm excitation/590 nm emission) to quantify total cholesterol content in media of Schwann cells 48 hours after siRNA transfection (B) without and (C) with ER stress induction by addition of 0, 2, 10, or 100 ng·mL^-1^ of tunicamycin. Total cholesterol content was normalized to total protein content in cells (mg protein) quantified using Bradford assay. All data presented in graphs are derived from *N* = 3 experimental repeats for each condition. Data expressed as mean ± SD. (B) One-way ANOVA and (A, C) two-way ANOVA with post-hoc Tukey’s test (*p<0.05, **p<0.01, ***p<0.001).

### Impact of Luman and ER stress on BDNF expression in Schwann cells

BDNF, a member of the neurotrophin family of growth factors is synthesized in both neurons and myelinating glial cells and is a key pro-myelinating molecule in the PNS, (Funakoshi et al., 1993; Karchewski et al., 2002; Zhang et al., 2000). It plays important roles in neuronal development, oligodendroglial proliferation, and remyelination of injured neurons. BDNF is initially produced as a precursor protein (proBDNF), which, in Schwann cells and neurons, undergoes folding and sorting into the regulatory secretory pathway (Lu et al., 2005) Following secretion, proBDNF molecules are cleaved by extracellular proteins into the mature form (mature BDNF – mBDNF). While the cleaved mBDNF is important for myelination and cell survival acting through the TrkB receptor tyrosine kinase, the predominant form of BDNF produced by Schwann cells and cultured neurons is the uncleaved proBDNF isoform (Mowla et al., 1999). proBDNF also exhibits neurotrophic activity by binding with high affinity to the p75^NTR^ neurotrophin receptor, which, when bound to the sortilin co-receptor, can initiate cell death-associated pathways (B. Lu et al., 2005; Teng et al., 2005). In addition to its roles in myelination and cell viability, BDNF initiates UPR signaling by activation of the IRE1 signaling arm and increases the splicing frequency of the *Xbp1* transcript in primary neuronal cultures, thus encoding the XBP1s transcription factor which in turn regulates BDNF expression in a positive feedback loop (Martínez et al., 2016; Saito et al., 2018) (Figure 5A). It has also been demonstrated that BDNF expression is heavily regulated by variations in the activation states of the PERK/ATF4 pathway in both neurons (J. Liu et al., 2018) and Schwann cells (B. Wang et al., 2017) in a TrkB-dependent manner. These previously described roles of BDNF in both the IRE1 and PERK arms of the UPR, as well as its well-established role in myelination, made us seek to determine the effect of Luman on BDNF expression and processing in the context of ER stress.

**Figure 5.**
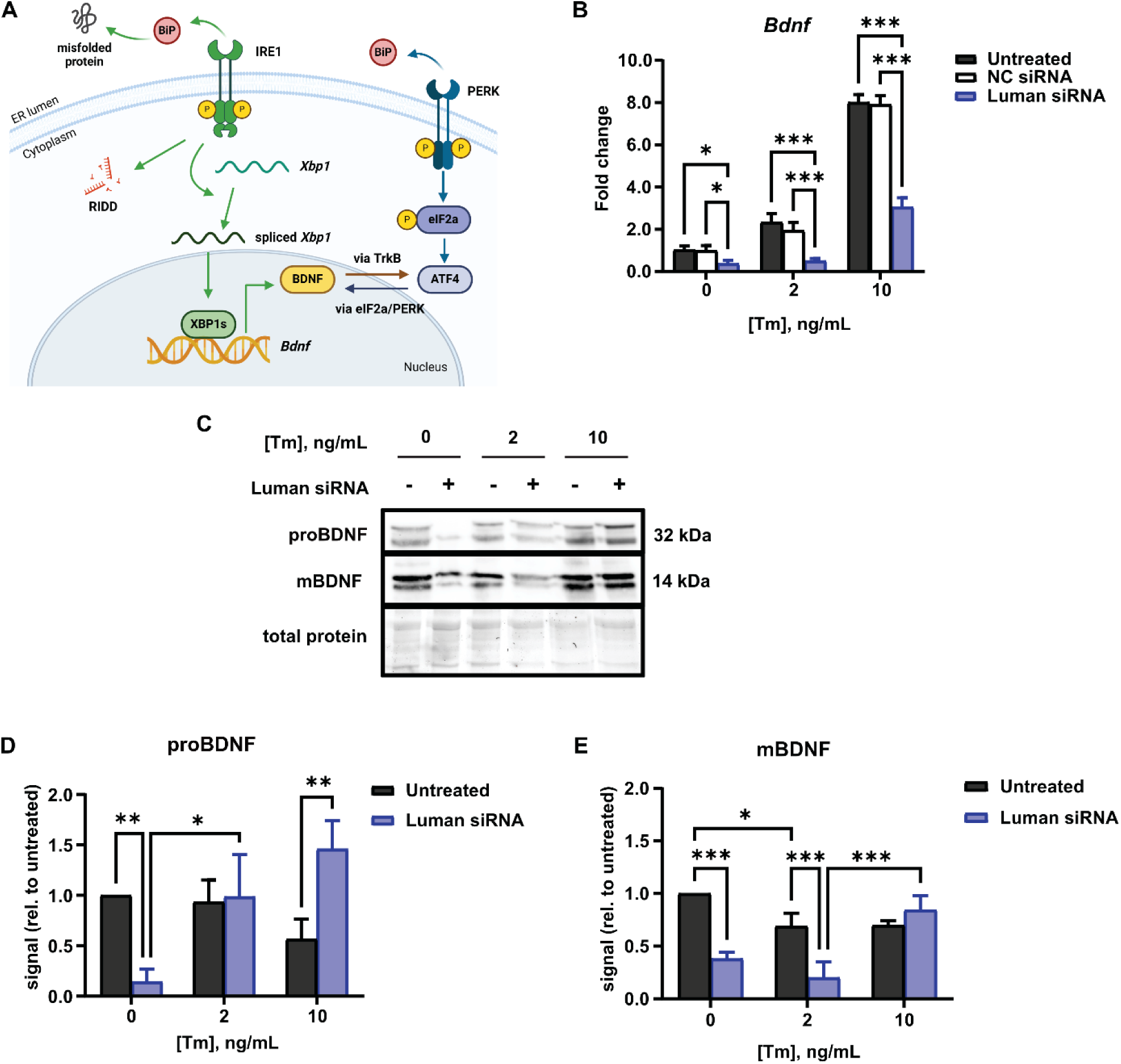
Brain-derived neurotrophic factor (BDNF) production and processing is directly influenced by varying Luman and ER stress levels in Schwann cells. (A) BDNF protein production is regulated by both spliced XBP1-mediated transcriptional regulation through the IRE1 arm and ATF4 signaling through the PERK arm of the unfolded protein response. Image created using BioRender. (B) Quantification of fold change by RT-qPCR of *Bdnf* transcript levels in Schwann cells 48 hours after siRNA transfection with or without ER stress induction by addition of 0, 2, or 10 ng·mL^-1^ of tunicamycin. (C) Representative immunoblots of protein isolated from Schwann cells 48 hours after siRNA transfection with or without ER stress induction by addition of 0, 2, or 10 ng·mL^-1^ of tunicamycin and probed for unprocessed proBDNF (prBDNF) at 32 kDa and mature BDNF (mBDNF) at 14 kDa. Densitometric analysis of band intensities was performed and data were normalized to total protein levels in each lane. Band intensities corresponding to (D) prBDNF and (E) mBDNF were then normalized to the mean band intensities of untreated groups. All data presented in graphs are derived from *N* = 3 experimental repeats for each condition. Data expressed as mean ± SD. Two-way ANOVA with post-hoc Tukey’s test (*p<0.05, **p<0.01, ***p<0.001).

Forty-eight hours following knockdown of Luman expression in Schwann cells, *Bdnf* transcript levels significantly decreased regardless of ER stress levels (Figure 5B). In addition, increasing ER stress to mild or moderate levels results in an overall increase of *Bdnf* levels in Schwann cells, which confirms what is known about how BDNF is a key target in transcriptional activation of the UPR in neuronal cell types. Although this is the case, it is important to know what isoform of BDNF is being produced following Luman knockdown and/or the induction of ER stress and this necessitates looking at variations in protein expression levels for the BDNF isoforms.

Luman knockdown in unstressed Schwann cells reduces proBDNF production when compared to the transfection control group (Figure 5C, D). However, at moderate ER stress levels, Schwann cells with reduced Luman levels show an increase in proBDNF levels compared to moderately stressed untransfected Schwann cells. This increase in proBDNF production following ER stress induction correlates with the observed increase in cell death at these conditions. Thus, moderate ER stress levels may reduce the capacity for the ER to properly package, process, and sort proBDNF produced by the cell towards the regulatory secretory pathway. Normally, the cell has the capacity to rectify any effects of this. However, Luman knockdown could contribute to the inability for the cells to mitigate any ER stress-associated cellular damage and could increase the probability of proBDNF binding to the p75^NTR^ receptor to initiate apoptosis.

The reduction in mBDNF levels following Luman knockdown in unstressed Schwann cells (Figure 5C, E) is also observed when analyzing proBDNF levels, and could indicate a possible indirect role for Luman in either transcriptional control of the *Bdnf* gene or synthesis of the proneurotrophin isoform itself. This pattern is maintained at mild ER stress levels. Levels of mBDNF also decrease upon induction of mild ER stress independent of Luman expression as shown in the decrease of mBDNF levels in the untransfected group (Figure 5C, E). This supports the idea that the increased number of proBDNF molecules produced in these conditions are either not being secreted or not being cleaved by extracellular proteases following secretion, or both. This observation correlates with the increased cell death observable following induction of ER stress in Schwann cells where Luman expression is knocked down (Figure 1B). Finally, increasing ER stress to moderate levels results in an increase of mBDNF protein in Schwann cells where Luman expression is knocked down compared to the untransfected and mildly stressed groups with reduced Luman expression (Figure 5C, E). As this increase is only observable in cells where Luman is knocked down, this could be attributed to possible cellular “leakiness” associated with an increased number of damaged cells following Luman knockdown contributing to proBDNF molecules (which increase in number under these conditions) being released into the extracellular environment to be cleaved by proteases, resulting in a rise in mBDNF levels without the need for undergoing sorting and packaging into the usual secretory pathway.

## DISCUSSION

This study reveals a novel role for Luman in regulating survival and myelination in Schwann cells in connection to its previously established role in the unfolded protein response and cholesterol biosynthesis pathways in sensory neurons associated with axon regeneration (Ying et al., 2014, 2015). Knockdown of Luman expression in primary Schwann cells significantly reduced cell viability in unstressed, mildly ER stressed, and moderately ER stressed cells 48 hours post-transfection with Luman siRNA. At the same time, there was a significant increase in apoptotic cell numbers at 48 hours and 72 hours post-transfection in unstressed and stressed Schwann cells with reduced Luman expression. Using myelination co-cultures with DRG sensory neurons, we observed an increase in MBP transcript and protein expression levels, as well as a general increase in myelin coverage over mature axons in co-cultures with Luman-overexpressing Schwann cells. Analysis of the effect of Luman on UPR mediators and targets inside and outside the context of ER stress, including BDNF, demonstrates an adaptive response that it can mount at lower levels of ER stress – a response that is generally different in the absence of ER stress in Schwann cells. Finally, Luman was also shown to have a significant influence on the amount of total cholesterol produced by Schwann cells and the expression levels of the major cholesterol precursor *Srebf1*. Collectively, these data reveal a possible cytoprotective role that Luman might play in Schwann cells to mitigate cellular damage from ER stress and promote an increased capacity for myelination. This response could directly facilitate the generation of a conducive environment for increased myelination by serving as a positive regulator for (i) Schwann cell survival; (ii) MBP expression; (iii) cholesterol synthesis; and (iv) BDNF synthesis and maturation, thereby allowing us to gain better insight on both the benefits of low-level stress on myelinating cells and the role of Luman in these responses.

### Luman is a key transcription factor for the regulation of Schwann cell survival

Increased cell death was observable in Schwann cell cultures 48 hours after knockdown of Luman expression in unstressed, mildly stressed, and moderately stressed conditions. At severe ER stress conditions, cell death resulting from the tenfold increase in concentration of the pharmacological ER stress inducer tunicamycin overtook any effect Luman had on the normal physiology of the cells. Tunicamycin induces ER stress and activates the UPR pathway by blocking *N-*linked glycosylation and arresting the cell cycle at the G1 phase. Previous studies have determined that Luman expression is not directly affected by addition of tunicamycin *in vitro* in astrocytes, breast cancer cell lines, and sensory neurons (DenBoer et al., 2005; Déry & LeBlanc, 2017; Ying et al., 2015). We have also confirmed this is the case in Schwann cells (Figure S2), allowing tunicamycin to be used for our experiments. The accompanying cell death observed following Luman expression knockdown in Schwann cells was also observable in the absence of ER stress, indicative of Luman’s role in their survival on a more fundamental level, perhaps in the context of either cell cycle integrity, cell proliferation, or secretory protein production – all of which have been evaluated and described in other cell types (Penney et al., 2018; Zhao et al., 2016). Previously, Luman was not shown to be essential in maintaining the survival of DRGs/sensory neurons, with Luman knockdown not having a significant effect on the cell numbers of injury-conditioned and intact sensory neurons compared to controls (Ying et al., 2015). However, in glioblastoma tumors, it has been demonstrated that Luman is essential as a tumor promoter in glioblastoma cells by promoting proliferation, invasion, and survival; knockdown of Luman resulted in increased cell death by apoptosis (Hu et al., 2019). This lends credence to the context-dependent role of Luman in cellular survival and the idea that Luman might be more essential to maintaining cell survival in actively proliferating glial cells compared to postmitotic neuronal cells. Luman has also been known to regulate cell survival through its interaction with the Herp protein in the context of ER stress. Upon ER stress-mediated UPR activation, Luman induces Herp expression by binding to the ERSE-II sequence in its promoter (Kokame et al., 2001; Liang et al., 2006). In the study by Liang et al. (2006), overexpression of the activated N-terminal form of Luman reduced tunicamycin-induced apoptosis in HeLa cells and increased the expression of Herp, suggesting that cellular tolerance for ER stress-induced apoptosis was positively regulated by Luman binding to Herp. Herp by itself has been known to similarly confer a cytoprotective effect to reduce ER stress-induced apoptosis (S. L. Chan et al., 2004; Hori et al., 2004). In Schwann cells particularly, a connection between low density lipoprotein receptor-related protein 1 (LRP1) and Luman in mediating Schwann cell function and survival during nerve injury has also been drawn by previous studies (Campana et al., 2006).

In the context of peripheral nerve repair, LRP1 has emerged as a possible mediator for normal Schwann cell function and increased pro-repair response to nerve injury. A potential role of Luman in Schwann cell survival similar or in connection to LRP1 in the context of the unfolded protein response could exist as LRP1 has also been demonstrated to reduce UPR-induced Schwann cell apoptosis through activation of the PI3K pathway (Mantuano et al., 2011), but this remains to be investigated.

### Luman positively regulates Schwann cell-mediated myelination of sensory neurons *in vitro*

After inducing myelination in the co-cultures, they were maintained until DIV 21, when the cultures were fixed and co-stained for MBP and NF200. MBP is an essential myelin protein associated with the initiation of myelination and formation of compact myelin in both the PNS and the CNS. Further, the use of MBP to detect and quantify myelin coverage has previously been established in the DRG/SC myelination models (Taveggia & Bolino, 2018). NF200 is a heavy-neurofilament marker for mature, large diameter axons – the kind that Schwann cells selectively myelinate (Feltri et al., 2016). To standardize myelin coverage across these types of axons in both co-cultures, MBP/NF200 ratios were computed by measuring the area covered by MBP fluorescence and dividing it over the area covered by NF200 fluorescence. Because naïve and not injury-conditioned DRG neurons are used in the co-cultures, NF200 values within the same focal point in the same culture are predicted to be constant and did so under the various conditions examined (Figure 2E). Calculating MBP/NF200 ratios to determine variations in myelin area coverage is an established protocol in other *in vitro* myelination cultures (Clark et al., 2017). The increase in MBP levels resulting from Luman overexpression indicates that (i) either a possible regulatory role of Luman in myelination through MBP transcription; or (ii) an indirect Luman-regulated pathway involved in myelination that consequently also increases MBP expression by increasing expression of myelination-adjacent factors and factors that encourage myelin formation. One of these possible indirect pathways involves the cholesterol biosynthesis pathway. MBP associates with the lipid membrane of neurons and the compaction and thermostability of the myelin formed as a result has been shown to be dependent on cholesterol content (Träger et al., 2020). The impact of Luman on cholesterol biosynthesis that we found could affect MBP production through this manner, or through Luman-mediated upregulation of BDNF and other pro-myelinating factors. BDNF has been shown to regulate early myelination through TrkB signaling, which affects myelination, synaptic plasticity, and cell survival (J. R. Chan et al., 2001; Zhang et al., 2000).

### Luman impacts the expression of UPR mediators under the IRE1 and PERK arms during ER stress in Schwann cells *in vitro*

ER stress is detected by three sensors: IRE1, PERK, and ATF6. Activation of the IRE1 pathway involves decoupling of the BiP molecule followed by dimerization and autophosphorylation of the IRE1 receptor, which initiates an adaptive cellular response to mitigate lower levels of ER stress. This response involves the decay of select mRNA templates through regulated IRE1-dependent decay (RIDD) as well as splicing of the *Xbp1* mRNA template into the transcriptional activator XBP1s (spliced XBP1 protein) (Siwecka et al., 2021). In contrast, under severe or sustained ER stress conditions, IRE1 activation regulates cell fate by transcriptional induction of genes under the proapoptotic c-Jun N-terminal kinase (JNK) pathway (Oyadomari & Mori, 2004; Urano et al., 2000). Activation of the PERK pathway, on the other hand, involves the phosphorylation of the eukaryotic translation initiation factor 2α (eIF2α), which inhibits general protein synthesis while promoting the translation of activating transcription factor-4 (ATF4), leading to the regulation of transcription of UPR targets such as CHOP (Oyadomari & Mori, 2004).

The marked increase of *Xbp1s* transcript levels in moderately ER-stressed control Schwann cells that is reduced following the knockdown of Luman levels (Figure 3B) validates the idea that increased ER stress in Schwann cells results in an adaptive Luman-regulated cellular response leading to a general increase in the production of *Xbp1s* and, presumably, an increased frequency of *Xbp1s*-regulated transcription of cellular response genes that mount an adaptive response against ER stress. At lower ER stress levels, ATF6 processing is activated alongside a more stringent RIDD protocol, and, as ER stress levels increase, the XBP1 splicing mechanism becomes activated via IRE1 phosphorylation and a broader RIDD protocol is established to degrade a larger mRNA pool. Higher ER stress levels may be overcome by allowing multiple XBP1/XBP1s alternative splicing rounds to be generated to mitigate it (Moore & Hollien, 2015; Yoshida et al., 2001). Though Luman knockdown reduced the levels of *Xbp1s* at moderate ER stress conditions, it was still significantly elevated in moderately stressed Schwann cells relative to unstressed and mildly stressed Schwann cells. This could reflect the shift from ATF6 activation to the activation of both ATF6 and IRE1 arms, resulting in not just an increase in *Xbp1* splicing rate, but also an increase in RIDD activity that ensures that *Xbp1* turnover is still higher than ER stress necessitates it to reach. This broadening of RIDD activity also ensures the degradation of pro-survival mRNAs, thus activating pro-death RIDD mechanisms and leading to increased cell death at these levels of ER stress (D. Han et al., 2009; Hassler et al., 2012). Continual production of *Xbp1s* at moderate ER stress levels is suggestive of the multiple XBP1/XBP1s splicing rounds hypothesized by Yoshida et al. (2001) to occur at higher ER stress levels.

It was not predicted that Luman knockdown would only significantly decrease CHOP protein levels under moderate ER stress conditions compared to untreated groups (Figure 3F, G). While, in general, CHOP is known to induce proapoptotic signaling pathways under ER stress conditions, its role in myelinating glial cells is more divergent. Previous studies have shown that persistent CHOP activation in Schwann cells resulting from severe or sustained ER stress can promote demyelination but is not directly linked to cell death (Gow & Wrabetz, 2009; Pennuto et al., 2008). Schwann cell dedifferentiation has been hypothesized to play a role in ameliorating ER stress and prevent cell death resulting from ER stress, similar to how the UPR system works upon activation, possibly in a c-Jun-mediated fashion (Gow & Wrabetz, 2009; Parkinson et al., 2008). Luman expression seems to regulate *Chop* transcription at adaptive/protective ER stress conditions, leading to reduced CHOP protein levels. Under adaptive ER stress conditions, the transcriptional induction of *Chop* is regulated by four elements in its promoter – two amino acid response elements (AARE1 and AARE2) that are activated by the PERK branch of the UPR and two ER stress response elements (ERSE1 and ERSE2) activated through both the IRE1 and/or the ATF6 pathway (Donati et al., 2006; Klymenko et al., 2019). As Luman is processed similarly to the ATF6 ER stress sensor and also plays a role in the IRE1 branch of the UPR, it could have an indirectly greater additive effect on *Chop* transcription by affecting the expression of UPR-associated transcription factors that could bind to these promoter regions, although this was not determined. Alternatively, since Luman expression affects multiple branches of the UPR, knocking down its expression might affect *Chop* transcription induction at multiple binding regions. This would occur at lower ER stress conditions as the cell is still mounting an adaptive response. However, once ER stress levels shift to severe, persistent PERK activation would override the effects of Luman knockdown on UPR inducers and CHOP levels in these groups should drastically elevate as well.

BDNF is both a known pro-myelinating factor and a UPR mediator and target. Upon activation of the IRE1 pathway, XBP1s binds to the promoter region of the BDNF gene to regulate its transcription (Martínez et al., 2016). In turn, BDNF is also able to positively regulate *Xbp1* splicing through protein kinase A (PKA)-dependent IRE1 activation. Increasing levels of ER stress result in a resolute increase in *Bdnf* transcript levels in Schwann cells regardless of Luman expression levels, lending credence to the idea of a positive feedback loop involved between the production of BDNF and the XBP1 spliced isoform protein. This preferentially activates the *Xbp1* splicing pathway rather than the pro-death RIDD pathway at low levels of ER stress. Knocking down Luman expression seems to also reduce overall *Bdnf* transcript levels in Schwann cells, supporting the existence of a possible inherent pro-survival and pro-myelinating role of Luman in Schwann cells. Increasing ER stress levels also increases *Bdnf* levels in Luman-deficient cells, but *Bdnf* mRNA levels are always lower in these cells compared to untreated controls. Once BDNF is translated, it undergoes proteolytic cleavage from an unprocessed proBDNF form to the cleaved mature BDNF. While it’s the mature BDNF homodimer that mainly populates the axo-glial space and binds to and activates the neurotrophin receptor TrkB resulting in pro-survival and pro-myelinating events, the proBDNF form can also occupy this space (M. Wang et al., 2021). ProBDNF shows a high affinity for p75^NTR^ and the protein sortilin to form the proBDNF/p75^NTR^/sortilin complex, which initiates pro-apoptotic pathways and neurite shrinkage (M. Wang et al., 2021). Due to their conflicting and contrasting functions, it was necessary when measuring BDNF protein expression levels to take into consideration the nature of the BDNF protein being analyzed. Induction of the UPR even at low levels was shown to increase levels of the proBDNF form when Luman is knocked down in Schwann cells. This negative correlation between proBDNF and Luman in the presence of ER stress could be the result of Luman conferring a protective role in Schwann cells during periods of high ER stress. Thus, a protective role appears to be lost when Luman is downregulated by siRNA, resulting in the increase of the pro-apoptotic proBDNF protein. On the other hand, mature BDNF levels were observed to be lower in uninduced and mild ER stress conditions in Luman-deficient Schwann cells. The pro-repair nature of this BDNF form alongside the fact that it decreases in expression level as Luman decreases alongside it lends confidence to the idea behind Luman being cytoprotective outside the UPR system, either in preparation for ER stress or to condition the cell for repair against damage. The connection between BDNF and Luman in myelinating glial cells is two-fold. Firstly, BDNF/TrkB signaling pathways normally activate different intracellular pathways, allowing flexibility of the Trk receptors in terms of conferring pro-survival functions to the cells. These include the PI3K/PKB pathway, the MAPK/Erk pathway, and the PLCγ/IP3 pathway (Bonni et al., 1999; Kaplan & Miller, 1997). BDNF signaling can also activate the transcription factor CREB and/or CREB-binding protein (CBP), which regulate neural plasticity, ER stress response, and cell survival (Bathina & Das, 2015; Kaplan & Miller, 1997). In addition to this, BDNF is also positively associated with cholesterol and triglyceride levels and general lipid metabolism (Bathina & Das, 2015; Pelleymounter et al., 1995). TrkB activation by BDNF has been shown to be essential for energy homeostasis, with fatty acids capable of normalizing BDNF levels and reducing oxidative damage, while BDNF itself has been shown to increase the rate of *de novo* cholesterol biosynthesis in cortical neurons in a TrkB-dependent manner (Suzuki et al., 2007).

### Luman expression positively regulates total cholesterol production at varying levels of ER stress in Schwann cells

The *Srebf1* gene codes for both SREBP1-a and SREBP1-c via variance in transcriptional initiation, and both regulate genes involved in cholesterol and fatty acid metabolism (Shimano, 2002). The role of SREBPs in myelination largely involves synthesis of cholesterol for initial myelin synthesis. While SREBPs are reliant on reduced cholesterol levels to become activated, *de novo* fatty acid synthesis through interaction with Fasn is also important for myelination and blocking this interaction can result in dysmyelination (Montani et al., 2018; Verheijen et al., 2003). Previously, Luman has been associated with *Srebf1* through the cholesterol biosynthesis and UPR pathways (Ying et al., 2015). Knockdown of Luman expression in 1-day crush-injured sensory neurons downregulated the expression of transcripts involved in cholesterol biosynthesis, such as *Insig1*, *Insig2*, and, notably, *Srebf1.* Free and total intracellular cholesterol levels were also depleted in neurons where Luman expression was knocked down. These effects were shown to be reversed by UPR induction with 2 ng/mL tunicamycin. In Luman-deficient sensory neurons, Ying et al. (2015) also observed further reduction of SREBP levels in response to exogenous cholesterol given to rescue regenerative axon growth, indicating a possible negative feedback loop with regards to SREBP regulation. *Srebf1* mRNA levels were observed to decrease in Schwann cells upon Luman knockdown, validating previous results observed regarding the connection between Luman expression and cholesterol synthesis and the levels of cholesterol-associated transcripts in sensory neurons. It is notable, however, that in Schwann cells, increasing levels of ER stress were associated with reduced *Srebf1* levels, indicating a possible difference in the way Schwann cells initiate cholesterol synthesis to sensory neurons. Schwann cells preferentially obtain their cholesterol for myelin production through *de novo* synthesis rather than obtaining it from circulating cholesterol stores or from astrocytes. Further, Schwann cells are transcriptionally inert to any extracellular changes in cholesterol levels (Pertusa et al., 2007). In connection to reduced *Srebf1* levels, Luman knockdown also reduced total cholesterol production in unstressed, mildly stressed, and moderately stressed Schwann cells. Results show that at severe ER stress levels, the effect of Luman knockdown is negated by sustained activation of the unfolded protein response, leading to continual *de novo* cholesterol production through other biochemical pathways in an attempt to mitigate high ER stress levels. Reduced total cholesterol levels following Luman knockdown indicates a direct link between the major cholesterol synthesis pathway SREBP and Luman in Schwann cells. While other pathways exist that contribute to cholesterol synthesis, it is also possible that Luman interacts with major mediators of these pathways outside the context of ER stress since the redundant mechanisms are known to exist in Schwann cells to replace cholesterol stores upon their depletion may also be inactive when Luman is depleted. One of these is the neuregulin-ErbB pathway, which is known to regulate cholesterol synthesis by stimulation of the PI3K pathway and, similar to the SREBP pathway, binding to the promoter of the key cholesterol synthesis enzyme 3-hydroxy-3-methylglutaryl-CoA reductase (HMGR) (Goldstein et al., 2006; Pertusa et al., 2007). Future analysis of whether Luman regulates this pathway could provide insight on how Luman affects cholesterol synthesis and Schwann cell myelination, as the neuregulin-ErbB pathway also regulates myelin production and Schwann cell maturation (Garratt et al., 2000).

In conclusion, we found in this study that Luman positively regulates Schwann cell survival, myelinating capacity, and cholesterol synthesis through the unfolded protein response. In addition to this, Luman was shown to confer a cytoprotective role through activation of the IRE1 pathway and increased expression and processing of BDNF under adaptive, low levels of ER stress. Understanding the role of ER stress-associated transcriptional elements such as Luman will shed light on how adaptive levels of cellular stress could be beneficial in the context of neural repair and demyelination.

## Data Availability

The data that support the findings of this study are available from the corresponding author upon reasonable request.

## Acknowledgements

We would like to thank R. Zhai and J. Johnston for their excellent technical assistance and Dr. A. Krishnan for the use of his lab’s inverted microscope to take the apoptosis images. This work was supported by Multiple Sclerosis Society of Canada grants #2362 & #920566 to VMKV; Canadian Institutes of Health Research grants #142238MOP-142238 & PJT-183666 to VMKV; and University of Saskatchewan College of Medicine Research Awards (CoMRAD) to VMKV. JMAN was supported by a University of Saskatchewan College of Medicine Graduate Research Award (CoMGRAD).

## References

Bathina, S., & Das, U. N. (2015). Brain-derived neurotrophic factor and its clinical implications. Archives of Medical Science : AMS, 11(6), 1164–1178. 10.5114/aoms.2015.56342

Bonni, A., Brunet, A., West, A. E., Datta, S. R., Takasu, M. A., & Greenberg, M. E. (1999). Cell survival promoted by the Ras-MAPK signaling pathway by transcription-dependent and - independent mechanisms. *Science (New York*, N.Y.), 286(5443), 1358–1362. 10.1126/science.286.5443.1358

Bouçanova, F., & Chrast, R. (2020). Metabolic interaction between Schwann cells and axons under physiological and disease conditions. Frontiers in Cellular Neuroscience, 14. 10.3389/fncel.2020.00148

Bracchi-Ricard, V., Nguyen, K., Ricci, D., Gaudette, B., Henao-Mejia, J., Brambilla, R., Martynyuk, T., Gidalevitz, T., Allman, D., Bethea, J. R., & Argon, Y. (2023). Increased activity of IRE1 improves the clinical presentation of EAE. FASEB Journal: Official Publication of the Federation of American Societies for Experimental Biology, 37(12), e23283. 10.1096/fj.202300769RR

Bunge, R. P. (1994). The role of the Schwann cell in trophic support and regeneration. Journal of Neurology, *242*(1 Suppl 1), S19-21. 10.1007/BF00939235

Campana, W. M., Li, X., Dragojlovic, N., Janes, J., Gaultier, A., & Gonias, S. L. (2006). The low-density lipoprotein receptor-related protein is a pro-survival receptor in Schwann cells: Possible implications in peripheral nerve injury. The Journal of Neuroscience: The Official Journal of the Society for Neuroscience, 26(43), 11197–11207. 10.1523/JNEUROSCI.2709-06.2006

Chan, J. R., Cosgaya, J. M., Wu, Y. J., & Shooter, E. M. (2001). Neurotrophins are key mediators of the myelination program in the peripheral nervous system. Proceedings of the National Academy of Sciences, 98(25), 14661–14668. 10.1073/pnas.251543398

Chan, S. L., Fu, W., Zhang, P., Cheng, A., Lee, J., Kokame, K., & Mattson, M. P. (2004). Herp stabilizes neuronal Ca2+ homeostasis and mitochondrial function during endoplasmic reticulum stress. The Journal of Biological Chemistry, 279(27), 28733–28743. 10.1074/jbc.M404272200

Clark, A. J., Kaller, M. S., Galino, J., Willison, H. J., Rinaldi, S., & Bennett, D. L. H. (2017). Co-cultures with stem cell-derived human sensory neurons reveal regulators of peripheral myelination. Brain, 140(4), 898–913. 10.1093/brain/awx012

Clayton, B. L. L., & Popko, B. (2016). Endoplasmic reticulum stress and the unfolded protein response in disorders of myelinating glia. Brain Research, 1648, 594–602. 10.1016/j.brainres.2016.03.046

DenBoer, L. M., Hardy-Smith, P. W., Hogan, M. R., Cockram, G. P., Audas, T. E., & Lu, R. (2005). Luman is capable of binding and activating transcription from the unfolded protein response element. Biochemical and Biophysical Research Communications, 331(1), 113–119. 10.1016/j.bbrc.2005.03.141

Déry, M.-A., & LeBlanc, A. C. (2017). Luman contributes to brefeldin A-induced prion protein gene expression by interacting with the ERSE26 element. Scientific Reports, 7(1), 42285. 10.1038/srep42285

Donati, G., Imbriano, C., & Mantovani, R. (2006). Dynamic recruitment of transcription factors and epigenetic changes on the ER stress response gene promoters. Nucleic Acids Research, 34(10), 3116–3127. 10.1093/nar/gkl304

Eldridge, C. F., Bunge, M. B., Bunge, R. P., & Wood, P. M. (1987). Differentiation of axon-related Schwann cells in vitro. I. Ascorbic acid regulates basal lamina assembly and myelin formation. Journal of Cell Biology, 105(2), 1023–1034. 10.1083/jcb.105.2.1023

Feltri, M. L., Poitelon, Y., & Previtali, S. C. (2016). How Schwann cells sort axons: New concepts. *The Neuroscientist: A Review Journal Bringing Neurobiology*, Neurology and Psychiatry, 22(3), 252–265. 10.1177/1073858415572361

Funakoshi, H., Frisén, J., Barbany, G., Timmusk, T., Zachrisson, O., Verge, V. M., & Persson, H. (1993). Differential expression of mRNAs for neurotrophins and their receptors after axotomy of the sciatic nerve. The Journal of Cell Biology, 123(2), 455–465. 10.1083/jcb.123.2.455

Garratt, A. N., Britsch, S., & Birchmeier, C. (2000). Neuregulin, a factor with many functions in the life of a schwann cell. *BioEssays: News and Reviews in Molecular*, Cellular and Developmental Biology, 22(11), 987–996. 10.1002/1521-1878(200011)22:11<987::AID-BIES5>3.0.CO;2-5

Goldstein, J. L., DeBose-Boyd, R. A., & Brown, M. S. (2006). Protein sensors for membrane sterols. Cell, 124(1), 35–46. 10.1016/j.cell.2005.12.022

Gow, A., & Wrabetz, L. (2009). CHOP and the endoplasmic reticulum stress response in myelinating glia. Current Opinion in Neurobiology, 19(5), 505–510. 10.1016/j.conb.2009.08.007

Hamanaka, R. B., Bobrovnikova-Marjon, E., Ji, X., Liebhaber, S. A., & Diehl, J. A. (2009). PERK-dependent regulation of IAP translation during ER stress. Oncogene, 28(6), 910–920. 10.1038/onc.2008.428

Han, C., Jin, L., Mei, Y., & Wu, M. (2013). Endoplasmic reticulum stress inhibits cell cycle progression via induction of p27 in melanoma cells. Cellular Signalling, 25(1), 144–149. 10.1016/j.cellsig.2012.09.023

Han, D., Lerner, A. G., Vande Walle, L., Upton, J.-P., Xu, W., Hagen, A., Backes, B. J., Oakes, S. A., & Papa, F. R. (2009). IRE1α kinase activation modes control alternate endoribonuclease outputs to determine divergent cell fates. Cell, 138(3), 562–575. 10.1016/j.cell.2009.07.017

Hasmatali, J. C. D., De Guzman, J., Zhai, R., Yang, L., McLean, N. A., Hutchinson, C., Johnston, J. M., Misra, V., & Verge, V. M. K. (2019). Axotomy induces phasic alterations in Luman/CREB3 expression and nuclear localization in injured and contralateral uninjured sensory neurons: Correlation with intrinsic axon growth capacity. Journal of Neuropathology & Experimental Neurology, 78(4), 348–364. 10.1093/jnen/nlz008

Hassler, J., Cao, S. S., & Kaufman, R. J. (2012). IRE1, a double-edged sword in pre-miRNA slicing and cell death. Developmental Cell, 23(5), 921–923. 10.1016/j.devcel.2012.10.025

Haze, K., Yoshida, H., Yanagi, H., Yura, T., & Mori, K. (1999). Mammalian transcription factor ATF6 is synthesized as a transmembrane protein and activated by proteolysis in response to endoplasmic reticulum stress. Molecular Biology of the Cell, 10(11), 3787–3799. 10.1091/mbc.10.11.3787

He, Y., Han, S., Li, H., Wu, Y., Jia, W., Chen, Z., Pan, Y., Cai, N., Wen, J., Li, G., Liang, J., Zhao, J., Liu, Q., Liang, H., Ding, Z., Huang, Z., & Zhang, B. (2024). CREB3 suppresses hepatocellular carcinoma progression by depressing AKT signaling through competitively binding with insulin receptor and transcriptionally activating RNA-binding motif protein 38. MedComm, 5(7), e633. 10.1002/mco2.633

Hiramatsu, N., Messah, C., Han, J., LaVail, M. M., Kaufman, R. J., & Lin, J. H. (2014). Translational and posttranslational regulation of XIAP by eIF2α and ATF4 promotes ER stress-induced cell death during the unfolded protein response. Molecular Biology of the Cell, 25(9), 1411–1420. 10.1091/mbc.E13-11-0664

Hori, O., Ichinoda, F., Yamaguchi, A., Tamatani, T., Taniguchi, M., Koyama, Y., Katayama, T., Tohyama, M., Stern, D. M., Ozawa, K., Kitao, Y., & Ogawa, S. (2004). Role of Herp in the endoplasmic reticulum stress response. Genes to Cells: Devoted to Molecular & Cellular Mechanisms, 9(5), 457–469. 10.1111/j.1356-9597.2004.00735.x

Hu, Y., Chu, L., Liu, J., Yu, L., Song, S., Yang, H., & Han, F. (2019). Knockdown of CREB3 activates endoplasmic reticulum stress and induces apoptosis in glioblastoma. Aging (Albany NY), 11(19), 8156–8168. 10.18632/aging.102310

Jang, S.-W., Kim, Y. S., Kim, Y. R., Sung, H. J., & Ko, J. (2007). Regulation of human LZIP expression by NF-kB and its involvement in monocyte cell migration Induced by Lkn-1. Journal of Biological Chemistry, 282(15), 11092–11100. 10.1074/jbc.M607962200

Kaewkhaw, R., Scutt, A. M., & Haycock, J. W. (2012). Integrated culture and purification of rat Schwann cells from freshly isolated adult tissue. Nature Protocols, 7(11), 1996–2004. 10.1038/nprot.2012.118

Kaplan, D. R., & Miller, F. D. (1997). Signal transduction by the neutrophin receptors. Current Opinion in Cell Biology, 9(2), 213–221. 10.1016/S0955-0674(97)80065-8

Karchewski, L. A., Gratto, K. A., Wetmore, C., & Verge, V. M. K. (2002). Dynamic patterns of BDNF expression in injured sensory neurons: Differential modulation by NGF and NT-3. The European Journal of Neuroscience, 16(8), 1449–1462. 10.1046/j.1460-9568.2002.02205.x

Kleitman, N., Wood, P. M., & Bunge, R. P. (1998). Tissue Culture Methods for the Study of Myelination. In Molecular and cellular approaches to neural development (2nd ed., pp. 337–377). Oxford University Press.

Klymenko, O., Huehn, M., Wilhelm, J., Wasnick, R., Shalashova, I., Ruppert, C., Henneke, I., Hezel, S., Guenther, K., Mahavadi, P., Samakovlis, C., Seeger, W., Guenther, A., & Korfei, M. (2019). Regulation and role of the ER stress transcription factor CHOP in alveolar epithelial type-II cells. Journal of Molecular Medicine (Berlin, Germany), 97(7), 973–990. 10.1007/s00109-019-01787-9

Kokame, K., Kato, H., & Miyata, T. (2001). Identification of ERSE-II, a new cis-acting element responsible for the ATF6-dependent mammalian unfolded protein response. The Journal of Biological Chemistry, 276(12), 9199–9205. 10.1074/jbc.M010486200

Liang, G., Audas, T. E., Li, Y., Cockram, G. P., Dean, J. D., Martyn, A. C., Kokame, K., & Lu, R. (2006). Luman/CREB3 induces transcription of the endoplasmic reticulum (ER) stress response protein Herp through an ER stress response element. Molecular and Cellular Biology, 26(21), 7999–8010. 10.1128/MCB.01046-06

Liu, J., Amar, F., Corona, C., So, R. W. L., Andrews, S. J., Nagy, P. L., Shelanski, M. L., & Greene, L. A. (2018). Brain-derived neurotrophic factor elevates activating transcription factor 4 (ATF4) in neurons and promotes ATF4-dependent induction of Sesn2. Frontiers in Molecular Neuroscience, 11. 10.3389/fnmol.2018.00062

Liu, Z., Lv, Y., Zhao, N., Guan, G., & Wang, J. (2015). Protein kinase R-like ER kinase and its role in endoplasmic reticulum stress-decided cell fate. Cell Death & Disease, 6(7), e1822–e1822. 10.1038/cddis.2015.183

Lu, B., Pang, P. T., & Woo, N. H. (2005). The yin and yang of neurotrophin action. Nature Reviews Neuroscience, 6(8), 603–614. 10.1038/nrn1726

Lu, R., Yang, P., O’Hare, P., & Misra, V. (1997). Luman, a new member of the CREB/ATF family, binds to herpes simplex VP16-associated host cellular factor. Molecular and Cellular Biology, 17(9), 5117–5126. 10.1128/MCB.17.9.5117

Mantuano, E., Henry, K., Yamauchi, T., Hiramatsu, N., Yamauchi, K., Orita, S., Takahashi, K., Lin, J. H., Gonias, S. L., & Campana, W. M. (2011). The unfolded protein response is a major mechanism by which LRP1 regulates Schwann cell survival after injury. The Journal of Neuroscience: The Official Journal of the Society for Neuroscience, 31(38), 13376–13385. 10.1523/JNEUROSCI.2850-11.2011

Martínez, G., Vidal, R. L., Mardones, P., Serrano, F. G., Ardiles, A. O., Wirth, C., Valdés, P., Thielen, P., Schneider, B. L., Kerr, B., Valdés, J. L., Palacios, A. G., Inestrosa, N. C., Glimcher, L. H., & Hetz, C. (2016). Regulation of memory formation by the transcription factor XBP1. Cell Reports, 14(6), 1382–1394. 10.1016/j.celrep.2016.01.028

Moncan, M., Mnich, K., Blomme, A., Almanza, A., Samali, A., & Gorman, A. M. (2021). Regulation of lipid metabolism by the unfolded protein response. Journal of Cellular and Molecular Medicine, 25(3), 1359–1370. 10.1111/jcmm.16255

Montani, L., Pereira, J. A., Norrmén, C., Pohl, H. B. F., Tinelli, E., Trötzmüller, M., Figlia, G., Dimas, P., von Niederhäusern, B., Schwager, R., Jessberger, S., Semenkovich, C. F., Köfeler, H. C., & Suter, U. (2018). De novo fatty acid synthesis by Schwann cells is essential for peripheral nervous system myelination. The Journal of Cell Biology, 217(4), 1353–1368. 10.1083/jcb.201706010

Moore, K., & Hollien, J. (2015). IRE1-mediated decay in mammalian cells relies on mRNA sequence, structure, and translational status. Molecular Biology of the Cell, 26(16), 2873–2884. 10.1091/mbc.E15-02-0074

Mowla, S. J., Pareek, S., Farhadi, H. F., Petrecca, K., Fawcett, J. P., Seidah, N. G., Morris, S. J., Sossin, W. S., & Murphy, R. A. (1999). Differential sorting of nerve growth factor and brain-derived neurotrophic factor in hippocampal neurons. Journal of Neuroscience, 19(6), 2069–2080. 10.1523/JNEUROSCI.19-06-02069.1999

Oñate, M., Catenaccio, A., Martínez, G., Armentano, D., Parsons, G., Kerr, B., Hetz, C., & Court, F. A. (2016). Activation of the unfolded protein response promotes axonal regeneration after peripheral nerve injury. Scientific Reports, 6, 21709. 10.1038/srep21709

Oyadomari, S., & Mori, M. (2004). Roles of CHOP/GADD153 in endoplasmic reticulum stress. Cell Death and Differentiation, 11(4), 381–389. 10.1038/sj.cdd.4401373

Parkinson, D. B., Bhaskaran, A., Arthur-Farraj, P., Noon, L. A., Woodhoo, A., Lloyd, A. C., Feltri, M. L., Wrabetz, L., Behrens, A., Mirsky, R., & Jessen, K. R. (2008). C-Jun is a negative regulator of myelination. The Journal of Cell Biology, 181(4), 625–637. 10.1083/jcb.200803013

Pelleymounter, M. A., Cullen, M. J., & Wellman, C. L. (1995). Characteristics of BDNF-induced weight loss. Experimental Neurology, 131(2), 229–238. 10.1016/0014-4886(95)90045-4

Penney, J., Taylor, T., MacLusky, N., & Lu, R. (2018). Luman/CREB3 plays a dual role in stress responses as a cofactor of the glucocorticoid receptor and a regulator of secretion. Frontiers in Molecular Neuroscience, 11, 352. 10.3389/fnmol.2018.00352

Pennuto, M., Tinelli, E., Malaguti, M., Del Carro, U., D’Antonio, M., Ron, D., Quattrini, A., Feltri, M. L., & Wrabetz, L. (2008). Ablation of the UPR-mediator CHOP restores motor function and reduces demyelination in Charcot-Marie-Tooth 1B mice. Neuron, 57(3), 393–405. 10.1016/j.neuron.2007.12.021

Pertusa, M., Morenilla-Palao, C., Carteron, C., Viana, F., & Cabedo, H. (2007). Transcriptional control of cholesterol biosynthesis in Schwann cells by axonal neuregulin 1. Journal of Biological Chemistry, 282(39), 28768–28778. 10.1074/jbc.M701878200

Saher, G., Quintes, S., Möbius, W., Wehr, M. C., Krämer-Albers, E.-M., Brügger, B., & Nave, K.-A. (2009). Cholesterol regulates the endoplasmic reticulum exit of the major membrane protein P0 required for peripheral myelin compaction. The Journal of Neuroscience, 29(19), 6094–6104. 10.1523/JNEUROSCI.0686-09.2009

Saher, G., & Simons, M. (2010). Cholesterol and myelin biogenesis. Sub-Cellular Biochemistry, 51, 489–508. 10.1007/978-90-481-8622-8_18

Saito, A., Cai, L., Matsuhisa, K., Ohtake, Y., Kaneko, M., Kanemoto, S., Asada, R., & Imaizumi, K. (2018). Neuronal activity-dependent local activation of dendritic unfolded protein response promotes expression of brain-derived neurotrophic factor in cell soma. Journal of Neurochemistry, 144(1), 35–49. 10.1111/jnc.14221

Schindelin, J., Arganda-Carreras, I., Frise, E., Kaynig, V., Longair, M., Pietzsch, T., Preibisch, S., Rueden, C., Saalfeld, S., Schmid, B., Tinevez, J.-Y., White, D. J., Hartenstein, V., Eliceiri, K., Tomancak, P., & Cardona, A. (2012). Fiji: An open-source platform for biological-image analysis. Nature Methods, 9(7), 676–682. 10.1038/nmeth.2019

Schirò, G., Iacono, S., Ragonese, P., Aridon, P., Salemi, G., & Balistreri, C. R. (2022). A brief overview on BDNF-Trk pathway in the nervous system: A potential biomarker or possible target in treatment of multiple sclerosis? Frontiers in Neurology, 13. 10.3389/fneur.2022.917527

Schröder, M., & Kaufman, R. J. (2005). ER stress and the unfolded protein response. Mutation Research, 569(1–2), 29–63. 10.1016/j.mrfmmm.2004.06.056

Shimano, H. (2002). Sterol regulatory element-binding protein family as global regulators of lipid synthetic genes in energy metabolism. Vitamins and Hormones, 65, 167–194. 10.1016/s0083-6729(02)65064-2

Shy, M. E., Tani, M., Shi, Y. J., Whyatt, S. A., Chbihi, T., Scherer, S. S., & Kamholz, J. (1995). An adenoviral vector can transfer lacZ expression into Schwann cells in culture and in sciatic nerve. Annals of Neurology, 38(3), 429–436. 10.1002/ana.410380313

Siwecka, N., Rozpędek-Kamińska, W., Wawrzynkiewicz, A., Pytel, D., Diehl, J. A., & Majsterek, I. (2021). The structure, activation and signaling of IRE1 and its role in determining cell fate. Biomedicines, 9(2), 156. 10.3390/biomedicines9020156

Suzuki, S., Kiyosue, K., Hazama, S., Ogura, A., Kashihara, M., Hara, T., Koshimizu, H., & Kojima, M. (2007). Brain-derived neurotrophic factor regulates cholesterol metabolism for synapse development. The Journal of Neuroscience, 27(24), 6417–6427. 10.1523/JNEUROSCI.0690-07.2007

Szegezdi, E., Logue, S. E., Gorman, A. M., & Samali, A. (2006). Mediators of endoplasmic reticulum stress-induced apoptosis. EMBO Reports, 7(9), 880–885. 10.1038/sj.embor.7400779

Taveggia, C., & Bolino, A. (2018). DRG neuron/Schwann cells myelinating cocultures. Methods in Molecular Biology (Clifton, N.J.), 1791, 115–129. 10.1007/978-1-4939-7862-5_9

Teng, H. K., Teng, K. K., Lee, R., Wright, S., Tevar, S., Almeida, R. D., Kermani, P., Torkin, R., Chen, Z.-Y., Lee, F. S., Kraemer, R. T., Nykjaer, A., & Hempstead, B. L. (2005). ProBDNF induces neuronal apoptosis via activation of a receptor complex of p75NTR and sortilin. The Journal of Neuroscience: The Official Journal of the Society for Neuroscience, 25(22), 5455–5463. 10.1523/JNEUROSCI.5123-04.2005

Träger, J., Widder, K., Kerth, A., Harauz, G., & Hinderberger, D. (2020). Effect of cholesterol and myelin basic protein (MBP) content on lipid monolayers mimicking the cytoplasmic membrane of myelin. Cells, 9(3), 529. 10.3390/cells9030529

Urano, F., Wang, X., Bertolotti, A., Zhang, Y., Chung, P., Harding, H. P., & Ron, D. (2000). Coupling of stress in the ER to activation of JNK protein kinases by transmembrane protein kinase IRE1. Science (New York, N.Y.), 287(5453), 664–666. 10.1126/science.287.5453.664

Urra, H., Dufey, E., Lisbona, F., Rojas-Rivera, D., & Hetz, C. (2013). When ER stress reaches a dead end. Biochimica et Biophysica Acta (BBA) - Molecular Cell Research, 1833(12), 3507–3517. 10.1016/j.bbamcr.2013.07.024

Verheijen, M. H. G., Chrast, R., Burrola, P., & Lemke, G. (2003). Local regulation of fat metabolism in peripheral nerves. Genes & Development, 17(19), 2450–2464. 10.1101/gad.1116203

Walter, P., & Ron, D. (2011). The unfolded protein response: From stress pathway to homeostatic regulation. Science (New York, N.Y.), 334(6059), 1081–1086. 10.1126/science.1209038

Wang, B., Ning, H., Reed-Maldonado, A. B., Zhou, J., Ruan, Y., Zhou, T., Wang, H. S., Oh, B. S., Banie, L., Lin, G., & Lue, T. F. (2017). Low-intensity extracorporeal shock wave therapy enhances brain-derived neurotrophic factor expression through PERK/ATF4 signaling pathway. International Journal of Molecular Sciences, 18(2), 433. 10.3390/ijms18020433

Wang, M., Xie, Y., & Qin, D. (2021). Proteolytic cleavage of proBDNF to mBDNF in neuropsychiatric and neurodegenerative diseases. Brain Research Bulletin, 166, 172–184. 10.1016/j.brainresbull.2020.11.005

Weil, M.-T., Möbius, W., Winkler, A., Ruhwedel, T., Wrzos, C., Romanelli, E., Bennett, J. L., Enz, L., Goebels, N., Nave, K.-A., Kerschensteiner, M., Schaeren-Wiemers, N., Stadelmann, C., & Simons, M. (2016). Loss of myelin basic protein function triggers myelin breakdown in models of demyelinating diseases. Cell Reports, 16(2), 314–322. 10.1016/j.celrep.2016.06.008

Wrabetz, L., D’Antonio, M., Pennuto, M., Dati, G., Tinelli, E., Fratta, P., Previtali, S., Imperiale, D., Zielasek, J., Toyka, K., Avila, R. L., Kirschner, D. A., Messing, A., Feltri, M. L., & Quattrini, A. (2006). Different intracellular pathomechanisms produce diverse Myelin Protein Zero neuropathies in transgenic mice. The Journal of Neuroscience: The Official Journal of the Society for Neuroscience, 26(8), 2358–2368. 10.1523/JNEUROSCI.3819-05.2006

Ying, Z., Misra, V., & Verge, V. M. K. (2014). Sensing nerve injury at the axonal ER: Activated Luman/CREB3 serves as a novel axonally synthesized retrograde regeneration signal. Proceedings of the National Academy of Sciences, 111(45), 16142–16147. 10.1073/pnas.1407462111

Ying, Z., Zhai, R., McLean, N. A., Johnston, J. M., Misra, V., & Verge, V. M. K. (2015). The unfolded protein response and cholesterol biosynthesis link Luman/CREB3 to regenerative axon growth in sensory neurons. The Journal of Neuroscience, 35(43), 14557–14570. 10.1523/JNEUROSCI.0012-15.2015

Yoshida, H., Matsui, T., Yamamoto, A., Okada, T., & Mori, K. (2001). XBP1 mRNA is induced by ATF6 and spliced by IRE1 in response to ER stress to produce a highly active transcription factor. Cell, 107(7), 881–891. 10.1016/s0092-8674(01)00611-0

Zhang, J. Y., Luo, X. G., Xian, C. J., Liu, Z. H., & Zhou, X. F. (2000). Endogenous BDNF is required for myelination and regeneration of injured sciatic nerve in rodents. The European Journal of Neuroscience, 12(12), 4171–4180.

Zhao, F., Liu, H., Wang, N., Yu, L., Wang, A., Yi, Y., & Jin, Y. (2020). Exploring the role of Luman/CREB3 in regulating decidualization of mice endometrial stromal cells by comparative transcriptomics. BMC Genomics, 21(1), 103. 10.1186/s12864-020-6515-2

Zhao, F., Wang, N., Yi, Y., Lin, P., Tang, K., Wang, A., & Jin, Y. (2016). Knockdown of CREB3/Luman by shRNA in mouse granulosa cells results in decreased estradiol and progesterone synthesis and promotes cell proliferation. PLOS ONE, 11(12), e0168246. 10.1371/journal.pone.0168246

