## Supplemental Figures 1-3 for "The ER stress transcription factor Luman/CREB3 is a novel regulator of Schwann cell survival and myelinating capacity through the activation of the unfolded protein response and cholesterol biosynthesis pathways"

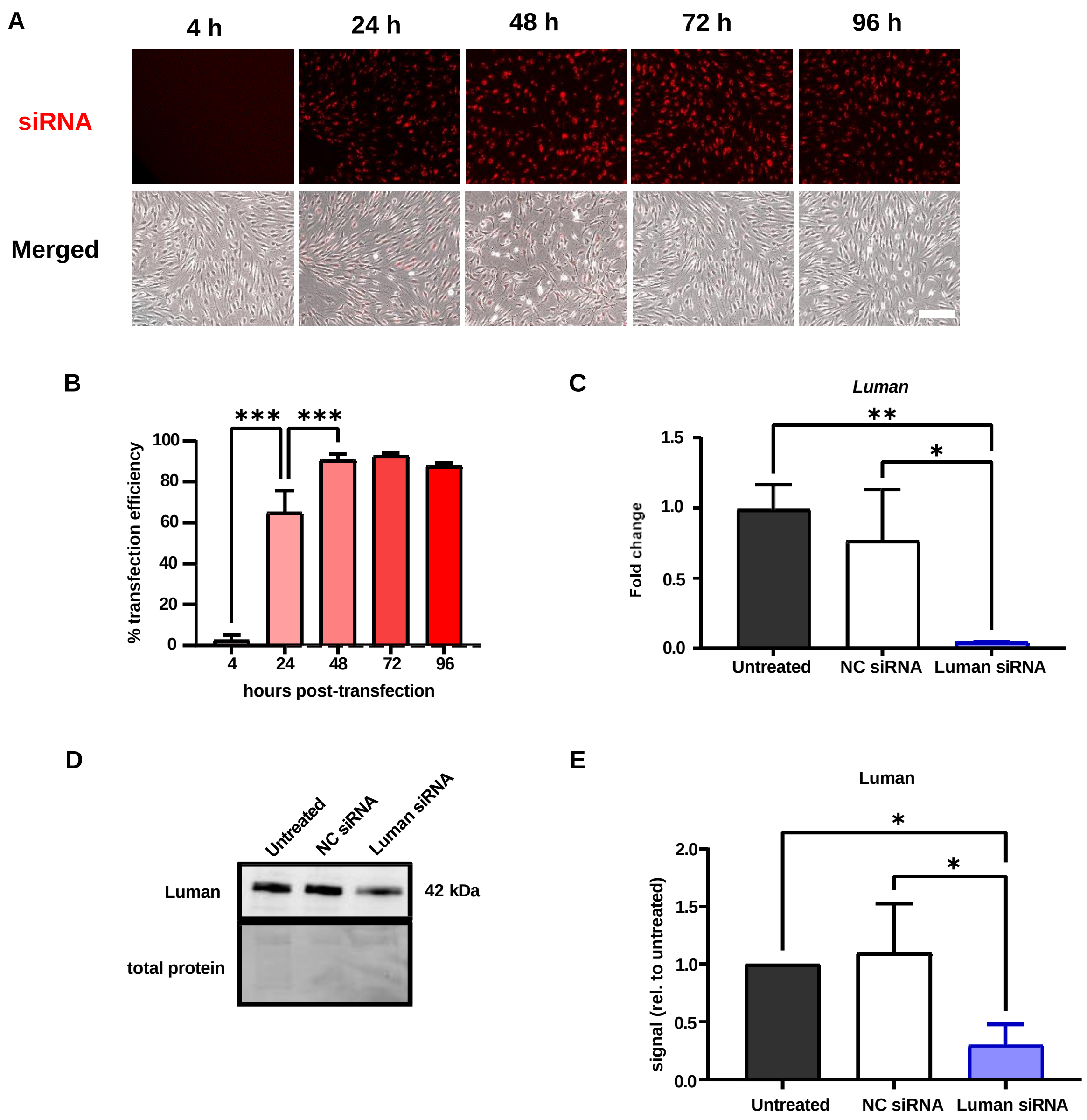


**Figure S1. Luman siRNA constructs and transfection protocols were validated and efficacy was quantified.** (A) Representative fluorescent micrographs of Schwann cells after varying timepoints (4 hours, 24 hours, 48 hours, 72 hours, and 96 hours post-transfection with 10 nM TYE 563-conjugated transfection control) showing positive transfectants (red).

Images in the red channel were merged with brightfield images. Scale bar = 100 μm. (B) Transfection efficiency was calculated by quantifying positive transfectants over total number of cells after each timepoint. (C) Quantification of fold change by RT-qPCR of *Luman* transcript levels in Schwann cells 48 hours after siRNA transfection. (D) Representative immunoblot of protein isolated from Schwann cells 48 hours after siRNA transfection and probed for Luman (42 kDa). Densitometric analysis of band intensities was performed and data were normalized to total protein levels in each lane. (E) Band intensities from Luman-probed immunoblots were normalized to the mean band intensities in untreated groups. All data presented in graphs are derived from $N = 3$ experimental repeats for each condition. Data expressed as mean ± SD. One-way ANOVA with post-hoc Tukey's test (*$p<0.05$, **$p<0.01$, ***$p<0.001$).

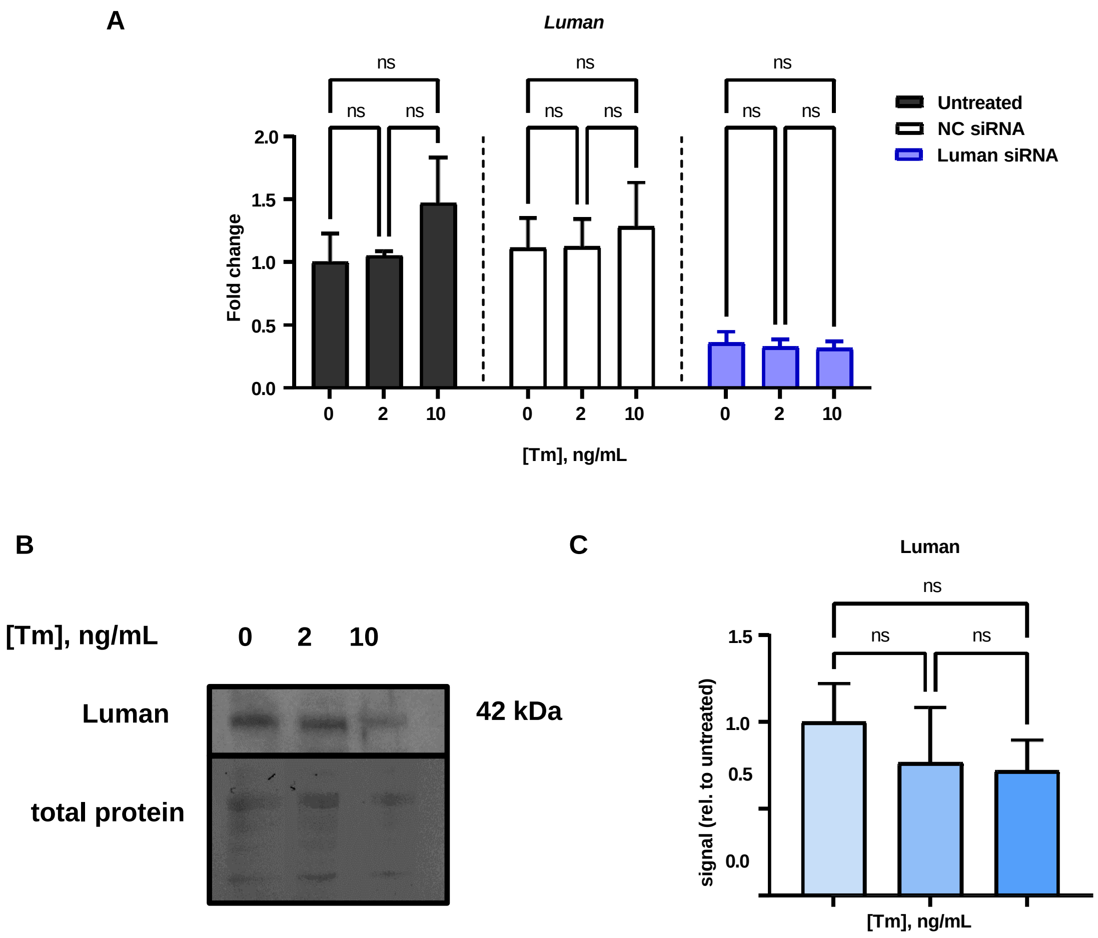


**Figure S2. Luman expression is not significantly affected by tunicamycin-induced ER stress.** (A) Quantification of fold change using RT-qPCR of *Luman* transcript levels in Schwann cells 48 hours after siRNA transfection with or without ER stress induction by addition of 0, 2, or 10 ng·mL$^{-1}$ of tunicamycin. Comparisons were made among different induction groups within the same transfection groups to determine significant differences. (B) Representative immunoblot of protein isolated from Schwann cells 48 hours after ER stress induction using 0, 2, or 10 ng·mL$^{-1}$ of tunicamycin and probed for Luman (42 kDa). Densitometric analysis of band intensities was performed and data were normalized to total protein levels in each lane. (C) Band intensities from Luman-probed immunoblots were then normalized to the mean band intensities of untreated groups. All data presented in graphs are derived from $N = 3$ experimental repeats for each condition. Data expressed as mean ± SD. One-way ANOVA with post-hoc Tukey's test (ns = no significant difference).

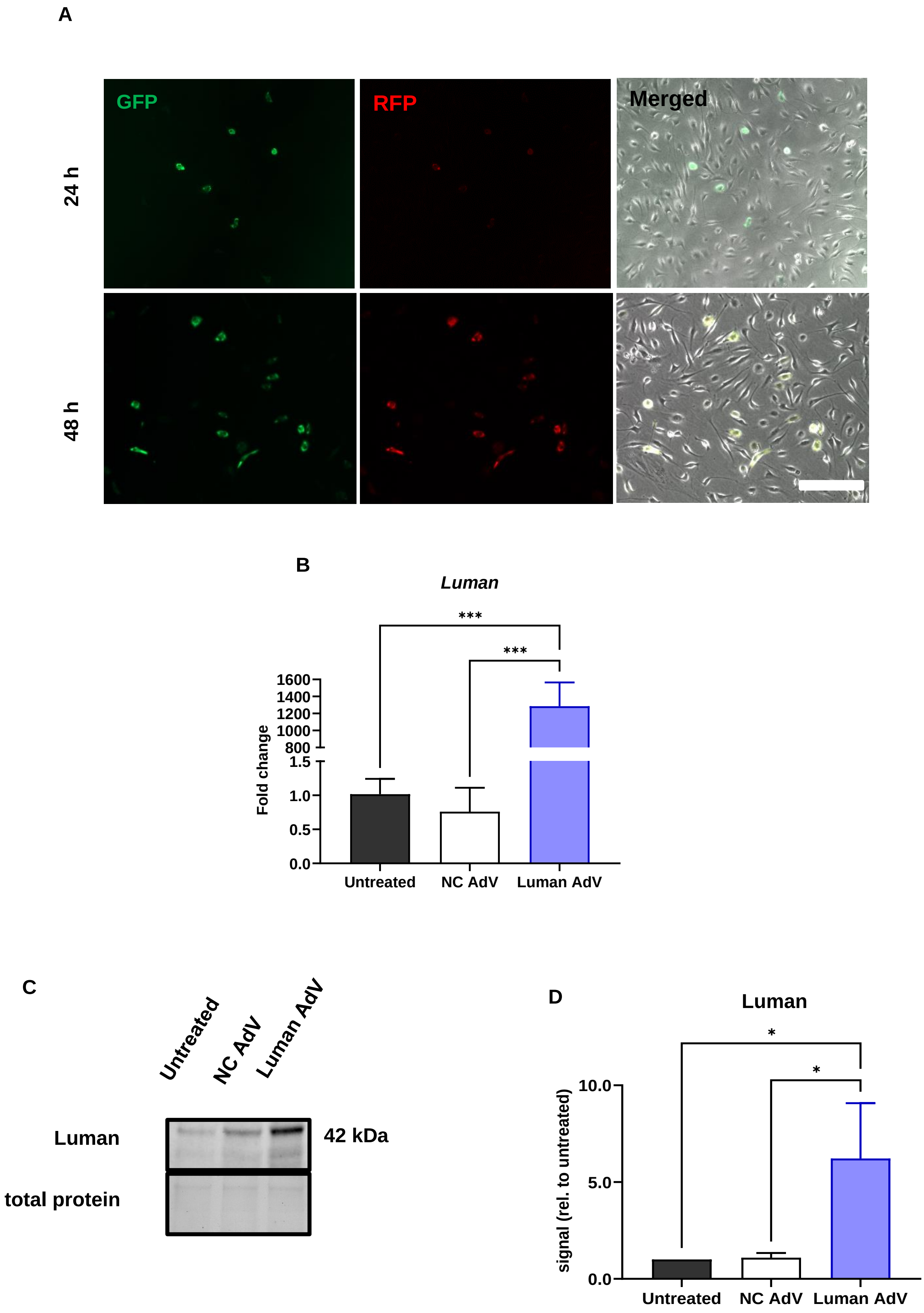

A
GFP
RFP
Merged
24 h
48 h
B
Luman
***
***
Fold change
1600
1400
1200
1000
800
1.5
1.0
0.5
0.0
Untreated
NC AdV
Luman AdV
C
Untreated
NC AdV
Luman AdV
Luman
42 kDa
total protein
D
Luman
*
*
signal (rel. to untreated)
10.0
5.0
0.0
Untreated
NC AdV
Luman AdV

**Figure S3. Luman adenovirus constructs and transduction protocols were validated and efficacy was quantified.** (A) Representative fluorescent micrographs of Schwann cells 24 hours and 48 hours post-transduction with AdV expressing GFP- (green) and RFP-conjugated (red) Luman sequence merged with brightfield images. Scale bar = 100 µm. (B) Quantification of fold change using RT-qPCR of *Luman* transcript levels in Schwann cells 48 hours after AdV transduction. (C) Representative immunoblot of protein isolated from Schwann cells 48 hours after AdV transduction and probed for Luman (42 kDa). Densitometric analysis of band intensities was performed and data were normalized to total protein levels in each lane. (D) Band intensities from Luman-probed immunoblots were then normalized to the mean band intensities in untreated groups. All data presented in graphs are derived from $N$ = 3 experimental repeats for each condition. Data expressed as mean ± SD. One-way ANOVA with post-hoc Tukey's test (*$p<0.05$, ***$p<0.001$).
